# BCG-induced reprogramming of monocyte/macrophage populations enhances lung antitumor immunity in mice

**DOI:** 10.1101/2025.07.25.666828

**Authors:** Eduardo Moreo, Miguel Araujo-Voces, Luna Minute, Laura Bravo-Robles, Ana Jiménez, Santiago Uranga, Ana Belén Gómez, Carlos Martín, Carlos del Fresno, Nacho Aguiló

## Abstract

The tumor microenvironment (TME) significantly influences antitumor immunity, with monocytes and macrophages playing pivotal roles both in pro- and anti-tumoral functions. Tumor-associated macrophages (TAMs) often adopt immunosuppressive phenotypes that promote tumor progression by inhibiting cytotoxic T and NK cells. This study investigates the antitumor mechanisms of intravenous (IV) Bacillus Calmette-Guérin (BCG) in a B16-F10 lung melanoma mouse model, focusing on its impact on monocyte/macrophage populations.

Single-cell RNA sequencing revealed that IV BCG reprograms tumor-associated monocyte-derived macrophages (mo-macs), shifting them from immunosuppressive to pro-inflammatory phenotypes enriched in interferon-response signatures. BCG treatment increased the recruitment of classical (Mon Iigp1) and non-classical (Mon Fcgr4) monocytes, which exhibited enhanced antigen presentation and pro-inflammatory cytokine production, while reducing immunosuppressive subsets prevalent in untreated controls. These BCG-induced mo-macs established robust interactions with NK and T cells, promoting their activation and enhancing cytotoxic function, as validated by functional assays. Notably, transfer of BCG reprogrammed bone marrow progenitors into naïve recipients elicited a sustained generation of immunostimulatory mo-macs that enhanced NK and T cell responses upon tumor challenge,

These findings highlight IV BCG’s potential as a cancer immunotherapy that targets the myeloid compartment to foster a pro-inflammatory TME, offering durable antitumor immunity by engaging both innate and adaptive immune responses.

## Introduction

The tumor microenvironment (TME) plays a central role in orchestrating antitumor immune responses. Among its diverse cellular constituents, monocytes and macrophages are key regulators of the tumor’s inflammatory landscape. These cells display remarkable functional plasticity, ranging from pro-inflammatory, antitumor phenotypes to immunosuppressive subsets that support tumor growth and progression^1^. Tumor-associated macrophages (TAMs) frequently adopt an immunosuppressive phenotype, employing a plethora of mechanisms to abrogate anti-tumor immunity. TAMs can secrete immunosuppressive cytokines such as IL-10 and TGF-β, which inhibit cytotoxic T cell activation, dampen NK cell cytotoxicity, and promote regulatory T cell (Treg) differentiation, thereby attenuating adaptive immune responses. TAMs also express high levels of immune checkpoint molecules which engage inhibitory receptors on T cells. Additionally, TAMs produce arginase-1 and indoleamine 2,3-dioxygenase (IDO), enzymes that deplete essential nutrients like arginine and tryptophan, impairing T cell proliferation and function. Through chemokine secretion (e.g., CCL22), TAMs recruit Tregs to the tumor site, further reinforcing an immunosuppressive milieu. Moreover, TAMs contribute to extracellular matrix remodeling and angiogenesis via VEGF and MMPs, hampering the access of immune effector cells to the tumor. These multifaceted immunosuppressive strategies position TAMs as key drivers of solid tumor progression and highlight their potential as therapeutic targets, owing to their inherent plasticity^2–6^.

NK cells play a pivotal role in antitumor immunity, particularly in suppressing metastasis. High NK cell infiltration is associated with reduced metastatic burden and improved survival outcomes across multiple cancer types^7^. NK cells recognize and eliminate malignant cells through activating receptors such as NKG2D, which detect stress-induced ligands (e.g., MICA/B, ULBPs) upregulated on tumor cells, and through reduced engagement of inhibitory receptors (e.g., KIRs, NKG2A) due to the frequent downregulation of MHC class I—an immune evasion strategy used by many tumor cells^8^. Reflecting their critical role in controlling metastasis, disseminating tumor cells often acquire specific phenotypes that enable evasion of NK cell recognition, such as increased expression of the NK cell-inhibitory ligand HLA-E^9^. Beyond their cytotoxic activity, NK cells also contribute to antitumor immunity within primary solid tumors by shaping the immune microenvironment. Notably, they can facilitate the recruitment of conventional type 1 dendritic cells (cDC1s)—key initiators of antitumor T cell responses—into tumors via the secretion of chemokines such as CCL5 and XCL1^10,11^. Importantly, NK cell antitumor activity can be inhibited by immunosuppresive TAMs^2^.

Bacillus Calmette-Guérin (BCG), a live attenuated vaccine originally developed for tuberculosis prevention, is also a first-line treatment for high-risk non-muscle invasive bladder cancer^12^. Intravenous (IV) administration of BCG provides superior protection against *Mycobacterium tuberculosis* challenge in macaques compared to intratracheal or subcutaneous routes, primarily by promoting robust immune cell recruitment to lung tissue^13–15^. Recently, we demonstrated that IV BCG could be harnessed for lung tumor treatment in mouse models ^16^. We showed that IV BCG activates an immune axis involving NK cells, cDC1s, and T cells. Specifically, BCG-stimulated NK cells promoted the recruitment and activation of cDC1s, facilitating the uptake of tumor-associated antigens and driving the development of tumor-specific CD8^+^ T cell responses. However, the mechanisms linking IV BCG administration to NK cell activation in the lung remained unclear, and the specific role of myeloid cell subsets in this process has not been fully explored yet. Notably, IV BCG is a potent inducer of trained immunity—a process in which innate immune cells undergo functional reprogramming at the level of hematopoietic progenitors, leading to enhanced responses upon subsequent challenges— which may be harnessed for cancer therapy^17–19^.

In this study, we sought to gain deeper insight into the antitumor mechanisms triggered by IV BCG in the lung. Using single-cell RNA sequencing (scRNAseq) in a mouse model of B16-F10 lung melanoma, we characterized the reprogramming of tumor-associated monocyte-derived macrophages (mo-macs) following IV BCG treatment and investigated their functional role in activating NK and T cells, as well as their contribution to the overall antitumor response.

## Results

### Intravenous BCG reshapes the lung tumor immune microenvironment

As we previously reported, therapeutic IV BCG administration reduces lung tumor burden in mice bearing various tumor types, including B16-F10 melanoma metastases. In the present study, we sought to characterize in an unbiased manner the composition and transcriptional states of immune cells within the tumor microenvironment (TME) following BCG treatment. To this end, we performed scRNAseq on sorted CD45⁺ cells from the lungs of B16-F10 tumor-bearing mice treated with either IV BCG or PBS as control. Samples were collected 20 days after tumor implantation—a time point at which the therapeutic effect of BCG is already detected (Fig. 1a).

**Figure 1.**
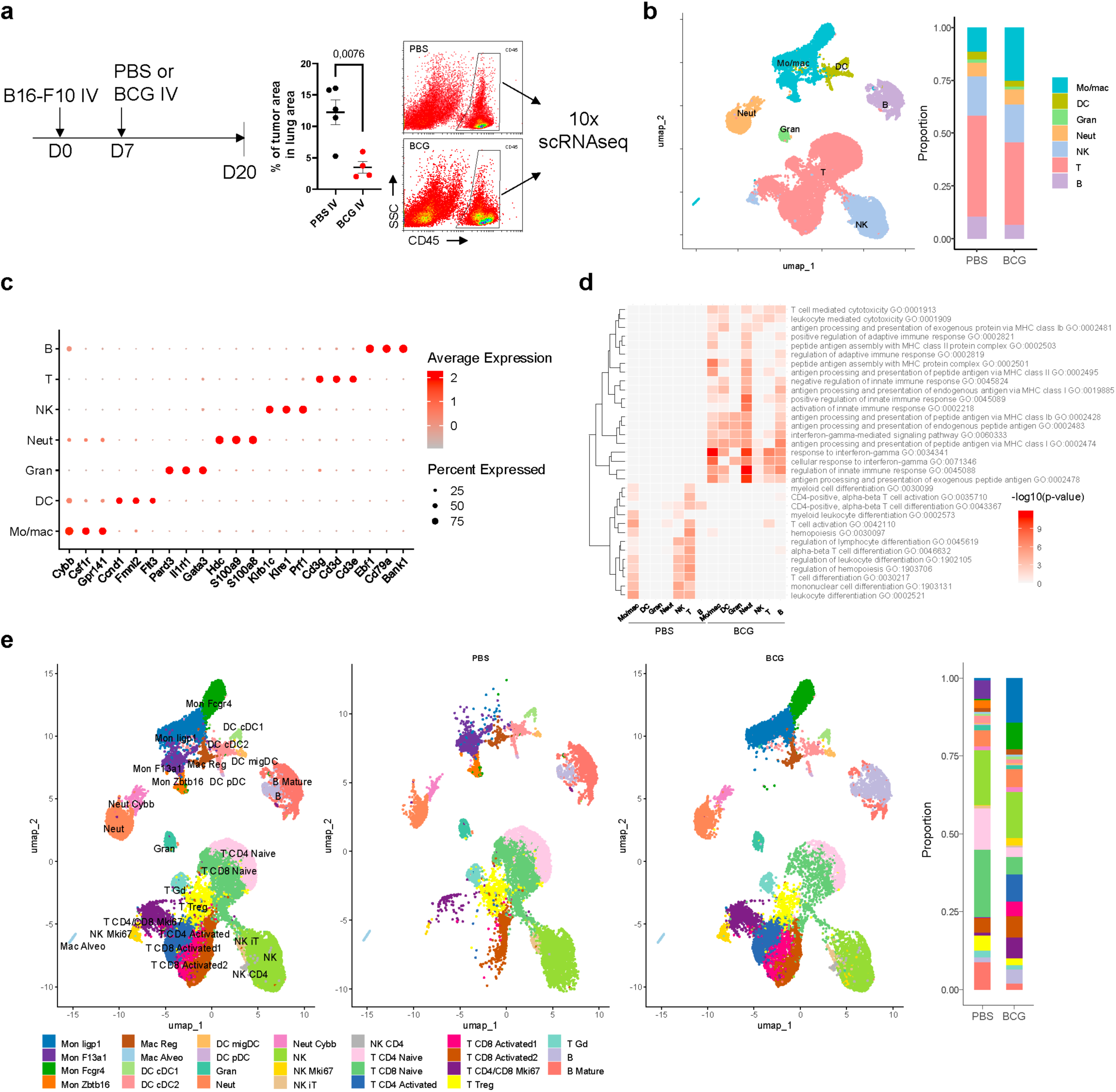
BCG treatment reshapes the immune landscape of B16-F10 lung tumors and enhances leukocyte activation. (**a**) Experimental design: mice were intravenously injected with B16F10 melanoma cells and treated with BCG or PBS on day 7. The endpoint was set at day 20 post-inoculation. Tumor burden was assessed, and CD45 cells were sorted for single-cell RNA sequencing using 10x Genomics technology. Lungs from five mice per condition were pooled and used for library construction. (**b**) UMAP and bar plot showing the proportions of major leukocyte clusters—monocytes/macrophages (mo/mac), dendritic cells (DCs), granulocytes (Gran), neutrophils (Neut), natural killer (NK) cells, T cells, and B cells—across BCG- and PBS-treated samples. (**c**) Dot plot displaying unbiased hub genes distinguishing general leukocyte populations. Dot size indicates the percentage of cells expressing each gene; color reflects average expression. (**d**) Gene ontology (GO) enrichment analysis of differentially expressed genes across general leukocyte populations. Hierarchical clustering of GO terms was performed using Euclidean distances based on adjusted p-values. (**e**) UMAP projection and bar plot illustrating subclustering, revealing 27 distinct leukocyte populations present in the experimental model.

Initial analysis identified seven clusters of immune cells: Monocytes/Macrophages (Mo/Mac), T cells, B cells, NK cells, neutrophils, granulocytes, and dendritic cells (Fig. 1b,c). GO analysis of differentially expressed (DE) genes among all clusters between groups pinpointed specific categories associated with BCG treatment, with an enrichment in inflammatory pathways related to type II IFN response, regulation of innate immune responses, and T cell-mediated cytotoxicity (Fig. 1d; Suppl. Tab. 1).

A deeper subclustering revealed twenty-seven cellular subsets, with substantial differences between PBS- and BCG-treated groups, particularly within T cell and mo-mac subsets (Fig. 1e, Suppl. Fig. 1a,b). Most T cells in the PBS condition clustered as T CD8 or T CD4 *naïve* cells, suggesting suboptimal activation following tumor challenge, whereas T cell activated subsets were more prevalent in BCG (Fig. 1e; Suppl. Fig. 1b,c). T CD8 activated subsets showed a higher cytotoxic gene signature compared to the control group, suggesting an enhanced cytotoxic potential induced by BCG (Suppl. Fig. 1d,e). Interestingly, the T CD4 activated cluster in the BCG condition expressed high levels of *Ifng* and *Csf1* transcripts, which could potentially be facilitating the activation, proliferation and survival of cells from the mo-mac lineage (Suppl. Fig. 1f)^20^. We next assessed exhaustion states using a curated set of T cell exhaustion markers (Suppl. Fig. 1g)^21,22^. Although no strong exhaustion phenotype was detected across conditions, T CD8 Activated2 and T CD4/CD8 Mki67 showed significantly lower exhaustion scores in BCG (Suppl. Fig. 1h). Lastly, proliferating T cells (*Mki67*, *Hist1h1b*, *Pclaf*) were more abundant following BCG treatment (Suppl. Fig. 1b,c).

When focusing on NK cells, we observed that this population clustered similarly under both conditions. However, a proliferative NK cell subset (characterized by *Mki67*, *Top2a*, and *Pclaf*) was markedly enriched in BCG-treated mice (Suppl. Fig. 1i-k). Furthermore, comparative analysis of NK clusters revealed a higher cytotoxic potential and enhanced functionality in the BCG-treated group (Suppl. Fig. 1l-o), in agreement with our previous data^16^. Regarding neutrophils and DCs, no major changes associated with treatment were found in the subclustering and functional signatures (Suppl. Fig. 2a-j). In the case of B cells, two clusters were identified: B (*Ly6a*, *Serpina3g*, *Agbl1*) and B Mature (*Ppm1h*, *Cd274*, *Trbc2*). UMAP projection, proportional analysis, and functional marker gene assessment revealed significant differences between both groups (Suppl. Fig. 2k-o).

### BCG reprograms tumor-associated monocyte/macrophage populations

We next examined mo-mac populations, which displayed profound alterations associated with BCG treatment (Fig. 2a, Suppl. Fig. 3a). Based on reported mo-mac PAN markers^23^, we identified two populations of classical monocytes, which express markers such us *Ccr2*, *Ly6c2* and *Cd14*: Mon Iigp1 (*Fgl2*, *Iigp1*, *Sell*) and Mon F13a1 (*F13a1*, *Ccl9*, *Stxbp6*); two of non-classical monocytes, characterized by downregulated *Ccr2* and *Cd14*, and expression of *Cx3cr1* and *Cd36*: Mon Fcgr4 (*Fcgr4*, *Smpdl3b*, *Cyfip2*) and Mon Zbtb16 (*Fabp4*, *Tgfbr3*, *Zbtb16*); and two populations of differentiated macrophages: alveolar macrophages (AM) (*Kcnip4*, *Atp6v0d2*, *Plet1*) and another enriched in *C1q* transcripts, which we termed Mac Reg (Fig. 2b; Suppl. Fig. 3a). Remarkably, our data revealed striking differences in the distribution of monocyte subpopulations between the two experimental conditions; Mon Iigp1 and Mon Fcgr4 populations were overrepresented in BCG-treated mice, whereas Mon F13a1 and Mon Zbtb16 were exclusively present in the untreated group (Fig. 2a; Suppl. Fig. 3a). Regarding differentiated macrophages, these populations appeared in both treatment conditions. Trajectory analysis suggested a directional progression from classical monocytes towards non-classical monocytes and Mac Reg (Suppl. Fig. 3b), likely reflecting the differentiation of blood monocytes to tumor-associated mature macrophages^3,24,25^.

**Figure 2.**
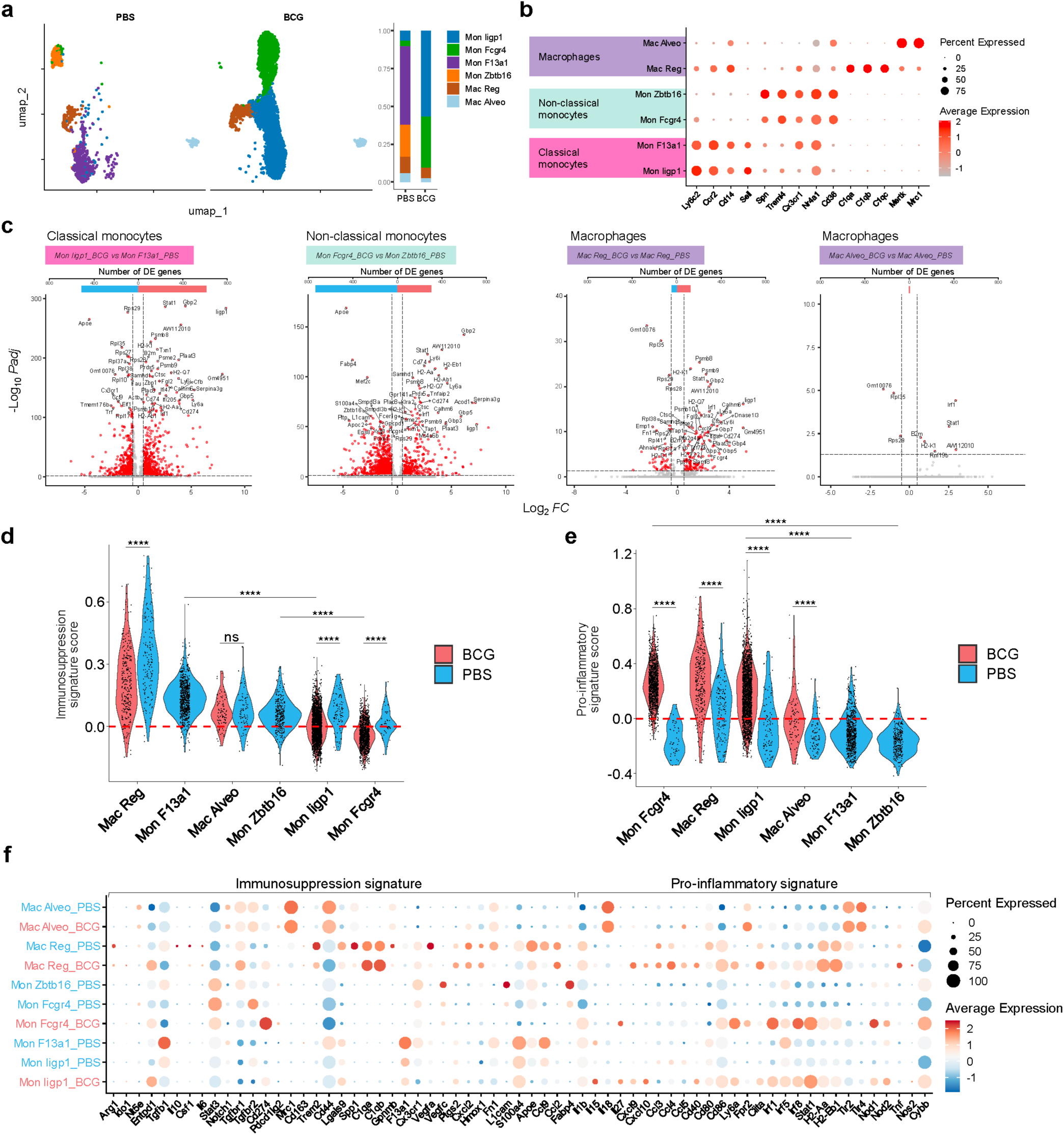
Transcriptional reprogramming of monocyte and macrophage subsets following BCG treatment. (**a**) UMAP plot and bar plot (proportions) displaying mo-macs subsets across PBS and BCG-treated samples. (**b**) Dot plot showing mo-macs PAN markers genes in classical, non-classical and differentiated macrophages. (**c**) Volcano plots showing DEGs between representative BCG- and PBS-associated subsets within classical monocytes, non-classical monocytes and macrophages. Specific comparatives are indicated below volcano titles. (**d**) Immunesuppression and (**e**) pro-inflammatoy signature analysis. Statistical analysis were performed using Wilcoxon tests, with p-value adjustment applied using the Bonferroni method. Significance levels are denoted as follows: ns = not significant (p ≥ 0.05), * = significant (p < 0.05), ** = highly significant (p < 0.01), *** = very highly significant (p < 0.001), and **** = extremely significant (p < 0.0001). (**f**) Dot Plot visualization of gene expression of curated gene lists. Dot size indicates the percent of cells expressing the gene and colors represent the average expression.

We next aimed to study differentially expressed (DE) genes between BCG- and PBS-specific mo-mac populations to identify gene features associated with treatment. First, we compared classical monocytes from both experimental conditions (Mon Iigp1 vs Mon F13a1), (Fig. 2b,c). Notably, key transcripts associated to immunosuppression were preferentially found in Mon F13a1 from PBS, with *Apoe* showing one of the highest fold changes (Fig. 2c; Suppl. Tab. 2)^26,27^. On the contrary, genes such as *Gbp2*, *Stat1, H2-Aa, H2-Aab1, Irf1, H2-K1, Cd74 and Cd274,* suggestive of response to type II IFN, were upregulated in Mon Iigp1 from the BCG condition (Fig. 2c), which was further confirmed by GO analysis (Suppl. Fig. 3c). Regarding non-classical monocytes, *Apoe* and *Fabp4*, previously involved in lipid accumulation in TAMs^28^, were enriched in Mon Zbtb16 from PBS, while *Stat1*, *Gbp2*, *H2-Aa*, *Acod1*, *Iigp1* and *H2-Ab1* were again enriched in Mon Fcgr4 from BCG (Fig. 2c). Finally, we directly compared Mac Reg and Mac Alveo from both conditions, given they were present in comparable proportions. In the case of Mac Reg, we found an enrichment of *Psmb8* and *Psmb9* in BCG, as well as other genes associated with IFNγ-mediated activation. PSMB8 and PSMB9 are proteins from the proteasome machinery, and their expression is associated with improved antigen presentation^29^. Consistent with this, analysis of The Kyoto Encyclopedia of Genes and Genomes (KEGG) phagosome gene signature further confirmed that this population exhibits the highest phagocytic activity among the tested subsets followed by classic monocytes in BCG (Suppl. Fig. 3d). Notably, Mac Alveo from BCG-treated mice showed only nine DE genes, most of which described as IFNγ-dependent (Fig. 2c).

In order to compare the immunosuppressive and inflammatory profiles of the different mo-mac subsets found in tumor-bearing lungs we defined respective specific signatures based on previous literature^2,6,23,30–32^. Mac Reg was the most immunosuppressive population, followed by Mon F13a1, Mac Alveo and Mon Zbtb16 (Fig. 2d). Remarkably, mo-mac subsets found in the PBS group displayed a higher immunosuppressive signature, whereas Mon Iigp1 and Mon Fcgr4, enriched in the BCG condition, appeared less immunosuppressive. In contrast, populations enriched in BCG showed the highest pro-inflammatory signature (Fig. 2e). Genes previously associated with immunosuppression, such as *Tgfb1, Apoe, Spp1, Vegfa* and *Gpnmb^26,33–36^*, were increased in the PBS condition across different mo-mac subsets (Fig. 2f). Interestingly, our data suggested a non-redundant distribution of some of these genes among the different populations.

For instance, Mac Reg highly expressed *Trem2, Spp1*, *Vegfa* and *Gpnmb*, whereas *Tgfb1* and *Ccl9* were enriched in the Mon F13a1 cells. In the case of Mon Zbtb16, higher expression of *Fabp4*, *L1cam* or *Vegfc* genes was noted (Fig. 2f). Comparison of the GO enriched in the different mo-mac subsets suggested certain specialization to execute distinct immunomodulatory functions (Suppl. Fig. 3c). This functional divergence was further supported by Reactome GSEA (Suppl. Fig. 3e), which revealed shared and subset-specific pathways, reinforcing the transcriptional and functional continuum between classical monocytes and differentiated macrophages. Another significant overexpressed gene in all Mon Iigp1, Mon Fcgr4 and Mac Reg BCG-associated populations was *Ly6a*, recently described as a marker of inflammatory monocyte-macrophages capable of presenting antigen and restimulating CD8^+^ T cells in the TME^37^. The different mo-mac subsets found in the BCG-condition also seemed to specialize in divergent inflammatory roles: Mon Iigp1 prominently expressed *Il15*, Mon Fcgr4 was associated with *Il27* and Mac Reg seemed to be specialized in the expression of *Cxcl9, Cxcl10, Ccl3, Ccl4*, *Ccl5* and *Tnf* (Fig. 2f). Overall, these results show that IV BCG treatment completely shifts the transcriptome of mo-mac populations in the lung, leading to the generation of two inflammatory monocyte subsets which are completely absent in PBS, and increasing the inflammatory potential of the mature macrophage subset.

Next, we characterized the cell-cell communication network between cell clusters using the CellChat algorithm, analyzing known Ligand-Receptor (L-R) pairs^38^. This approach measures the strength of potential signaling between cells, identifying Mon Iigp1 in the BCG condition as the most active cluster (Fig. 3a). We also identified T CD8 activated2 and NK cells as highly active clusters receiving signals in the BCG condition. Analyzing the directionality of the most differential interactions, we observed a clear association between Mon Iigp1 and NK cells and several effector T cells in the BCG condition, whereas in PBS, data suggested a strong signal influx from Mon F13a1 to NK cells and naïve T cells (Fig. 3b,c). In the case of Mac Reg, we found that they strongly interact with T cells and other populations of monocytes and macrophages, whereas less differentiated monocytes focused their interactions on NK cells (Fig. 3b,c). Overall, these data suggest that mo-mac populations establish strong interactions with both NK and T cells in the tumor-bearing lung, regardless of treatment condition.

**Figure 3.**
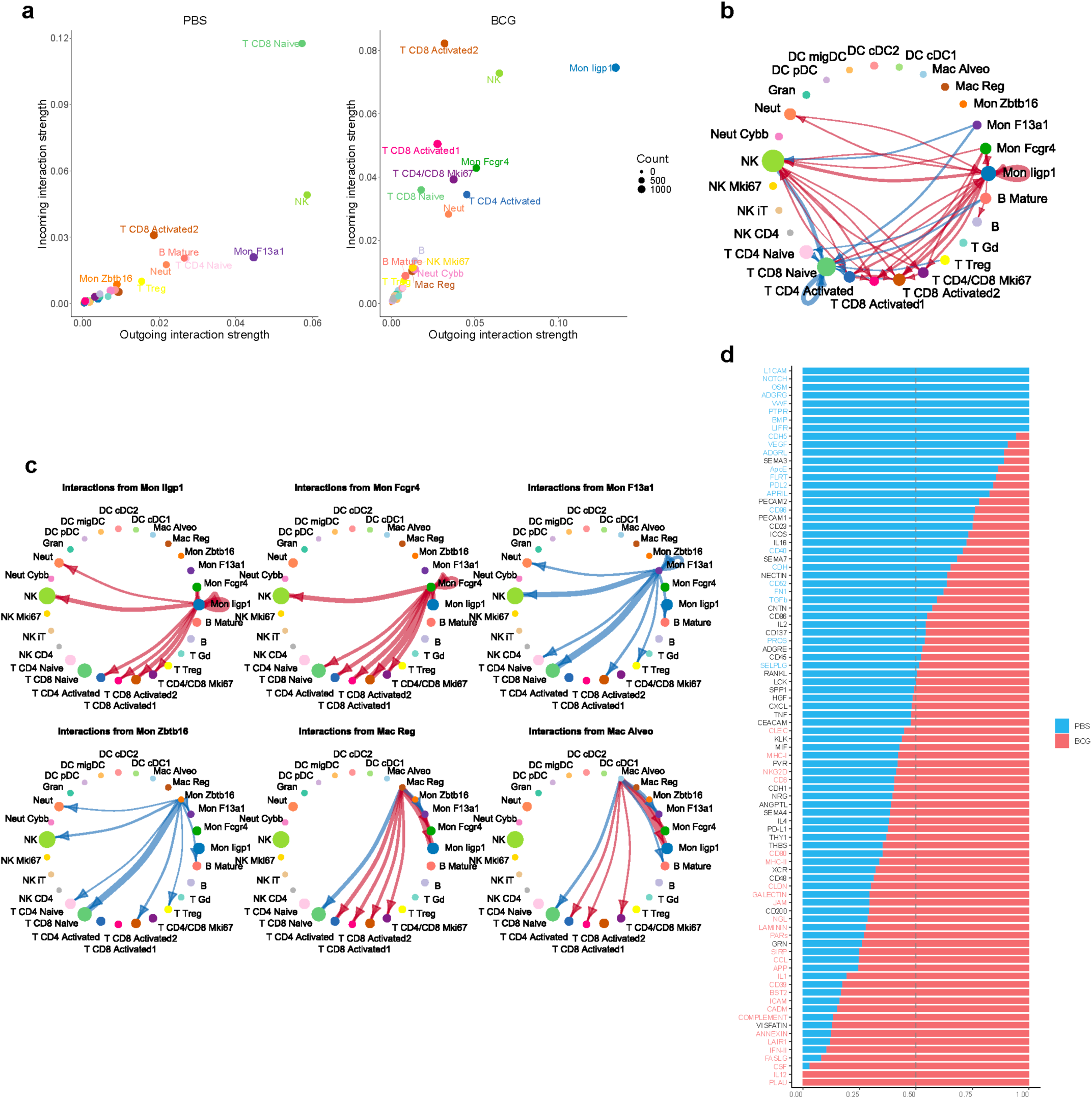
Cell-cell interaction patterns across cell types in BCG and PBS-treated samples. **(a)** Scatter plot showing outgoing (x-axis) and incoming (y-axis) interaction strengths across single-cell populations. Colors represent different cell populations, and dot size is proportional to the number of inferred links. All signaling pathways are included in this analysis. (**b**) Circle plot displaying the differential interaction strength in the cell-cell communication network. Only the top 5% of interactions are shown for both conditions. Line thickness represents the interaction strength, and arrows indicate directionality. (**c**) Circle plot highlighting the top 1% of differential interaction strengths. Each circle represents a specific monocyte-macrophage (mo-macs) subset as sender cells. (**d**) Bar chart illustrating the overall information flow in terms of signaling pathways. Colored pathways indicate those with significant differences based on a paired Wilcoxon test.

We then compared the overall information flow of L-R pairs, to identify the signaling pathways differentially expressed in each treatment condition. PBS-treated populations exhibited enriched pathways such as NOTCH, OSM, VEGF, ApoE or TGF-β, which have been previously associated with immune tolerance, tumor progression, and invasion (Fig. 3d; Suppl. Tab. 3)^33,39–43^. In contrast, BCG condition revealed enriched pathways associated with immune activation and cytotoxicity, such as IL12, MHC-I, NKG2D, CD80, MHC-II, CCL, APP, IFN-II, FASLG, and CSF^44,45^.

Altogether, our data indicate that IV BCG profoundly remodels the mo-mac compartment in the tumor-bearing lung. Ligand-receptor interaction analysis identified these BCG-induced mo-mac populations as a central signaling hub, with a clear directionality of communication toward NK and T lymphocytes and an enrichment in pathways associated with immune activation.

### Mo-macs are essential for the antitumoral effect of IV BCG

Next, we set out to validate scRNAseq results by flow cytometry and functionally assess the contribution of mo-macs to the BCG-induced antitumoral response. Tumor-bearing lung single cell suspensions were analyzed at day 20 post-tumor implantation, as in scRNAseq studies. Mo-macs were defined as CD11b⁺ SiglecF⁻ CD64⁺ cells by flow cytometry—a non-lung resident population that encompasses a spectrum of differentiation states from monocytes to macrophages in lung tissue. Higher numbers and frequencies of mo-macs were found in IV BCG-treated mice (Fig. 4a), as well as increased expression of MHC-II, PD-L1, STAT1, and CXCL9 (Fig. 4b,c), suggestive of enhanced IFNγ signaling in these cells, in agreement with scRNAseq data. On the contrary, tissue resident AMs (SiglecF^+^ CD64⁺ cells) were reduced both in total numbers and proportion following IV BCG treatment (Suppl. Fig. 4a). Furthermore, the enhanced activation status of mo-macs was not fully observed in AMs, with the exception of slightly increased CXCL9 and MHC-II (Suppl. Fig. 4b).

**Figure 4.**
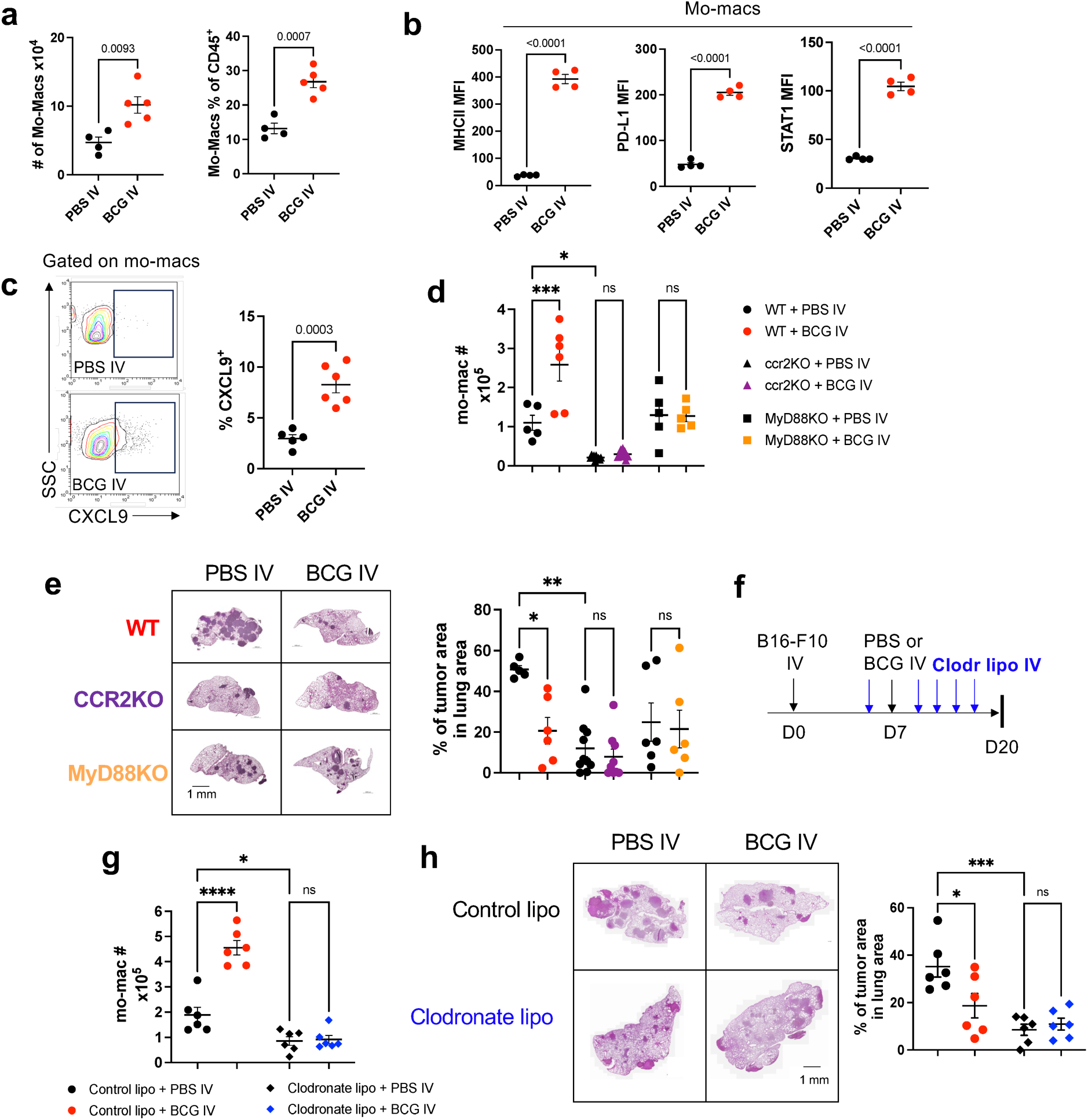
Effect of IV BCG on lung mo-mac populations. **(a)** Absolute numbers and % of mo-macs in the lungs of mice bearing B16-F10 lung tumors at day 20, analyzed by flow cytometry of single cell suspensions. **(b)** Surface expression of MHC-II and PD-L1 and intracellular expression of STAT1 in lung mo-macs. **(c)** Intracellular expression of CXCL9 in lung mo-macs. **(d)** Absolute numbers of mo-macs in the lungs of mice bearing B16-F10 lung tumors at day 20, analyzed by flow cytometry. **(e)** Representative H&E images of tumor-bearing lungs of PBS or BCG-treated mice at day 20 after inoculation of B16-F10 tumor cells, and quantification of the tumor area in lung cross-sections. Scale bars correspond to 1 mm in length. Data is pooled from 2 independent experiments **(f)** Schematic diagram showing treatment strategy for **(g-h) (g)** Absolute numbers and % of mo-macs in the lungs of mice bearing B16-F10 lung tumors at day 20, analyzed by flow cytometry of single cell suspensions. **(h)** Representative H&E images of tumor-bearing lungs of PBS or BCG-treated mice at day 20 after inoculation of B16-F10 tumor cells, and quantification of the tumor area in lung cross-sections. Scale bars correspond to 1 mm in length. Data is from 1 experiment. *P* values were calculated using two-tailed unpaired Student’s t test at a 95 % CI **(a,b,c)** or one-way ANOVA with Bonferroni multiple-comparison test **(d,e,g,h)**. Significance levels are denoted as follows: ns = not significant (p ≥ 0.05), * = significant (p < 0.05), ** = highly significant (p < 0.01), *** = very highly significant (p < 0.001)

Using a GFP-expressing BCG strain to monitor *in vivo* engulfing of BCG following IV administration, we identified that the vaccine was preferentially acquired by mo-macs, rather than AMs (Suppl. Fig. 4c). This result contrasted with the intranasal (IN) administration, through which BCG was mostly engulfed by AMs, in agreement with previous data^46^. Overall, these results suggest that BCG given by the IV route is preferentially taken up by mo-macs and contributes to their activation.

The higher numbers of mo-macs found in BCG-treated mice could be explained either by increased proliferation of tissue-resident mo-macs or recruitment from bone marrow-derived circulating monocytes. To discern these two possibilities, we made use of CCR2KO mice, in which monocyte migration to peripheral tissues is impaired. IV BCG was unable to increase lung mo-mac numbers in CCR2KO mice (Fig. 4d), evidencing that mo-macs were being mostly recruited to the tumor from circulating monocytes, via CCR2. Additionally, lack of CCR2 signaling impaired mo-mac recruitment also in untreated mice (Fig. 4d), showing that these cells are actively recruited to the lung during tumor growth.. We also assessed BCG in mice lacking MyD88, a crucial adaptor for innate sensing of pathogen-associated molecular patterns (PAMPs) and IL1beta receptor. Previous data indicate that these pathways are required to trigger BCG antitumoral responses^47^. In agreement with these results, our data revealed that BCG-driven mo-mac recruitment to the tumor-bearing lung was abrogated in mice lacking MyD88 (Fig. 4d).

Analysis of lung tumor burden showed that BCG antitumoral efficacy was impaired in both CCR2KO and MyD88KO mice (Fig. 4e), indicating the importance of innate sensing of BCG via MyD88 and subsequent CCR2-dependent mo-mac recruitment to the lung for the antitumoral response of IV BCG. Additionally, data from untreated CCR2KO mice revealed that recruitment of mo-macs via CCR2 facilitates tumor growth as recruited monocytes are a source of TAM (Fig. 4e), in agreement with previous data^48^.

To confirm these results with an alternative strategy, we depleted circulating monocytes by IV inoculation of clodronate liposomes (Fig. 4f), an approach previously used^49^. Clodronate liposome administration completely abrogated lung infiltration of mo-macs (Fig. 4g), as well as preventing BCG antitumoral efficacy (Fig. 4h). Altogether, our results show that IV BCG treatment triggers a CCR2 and MyD88-dependent influx of mo-macs which appear to be beneficial for antitumor immunity.

### BCG-activated mo-macs enhance NK and T cell function

Our scRNAseq data described a robust interaction between mo-macs and NK/T cells, both in untreated mice and in the context of BCG immunotherapy. In addition, our previous results indicated that IV BCG strongly activates lung NK and T cells *in vivo*. Therefore, we next assessed whether BCG-activated macrophages were mediating NK and T cell activation in the lung. We observed that both CCR2KO and MyD88KO impaired BCG-mediated NK cell activation, measured by IFNγ, Granzyme B and KLRG1 expression (Fig. 5a,b). A similar result was observed when IFNγ production by T cells was assessed, both for CD8+ and CD4+ subsets (Fig. 5c). Comparable results were obtained when mo-macs were depleted with clodronate liposomes (Suppl. Fig. 5a,b,c). Overall, these results indicate that mo-mac recruitment and activation driven by IV BCG crucially contribute to the enhancement of NK and T cell antitumoral function in lung tumors.

**Figure 5.**
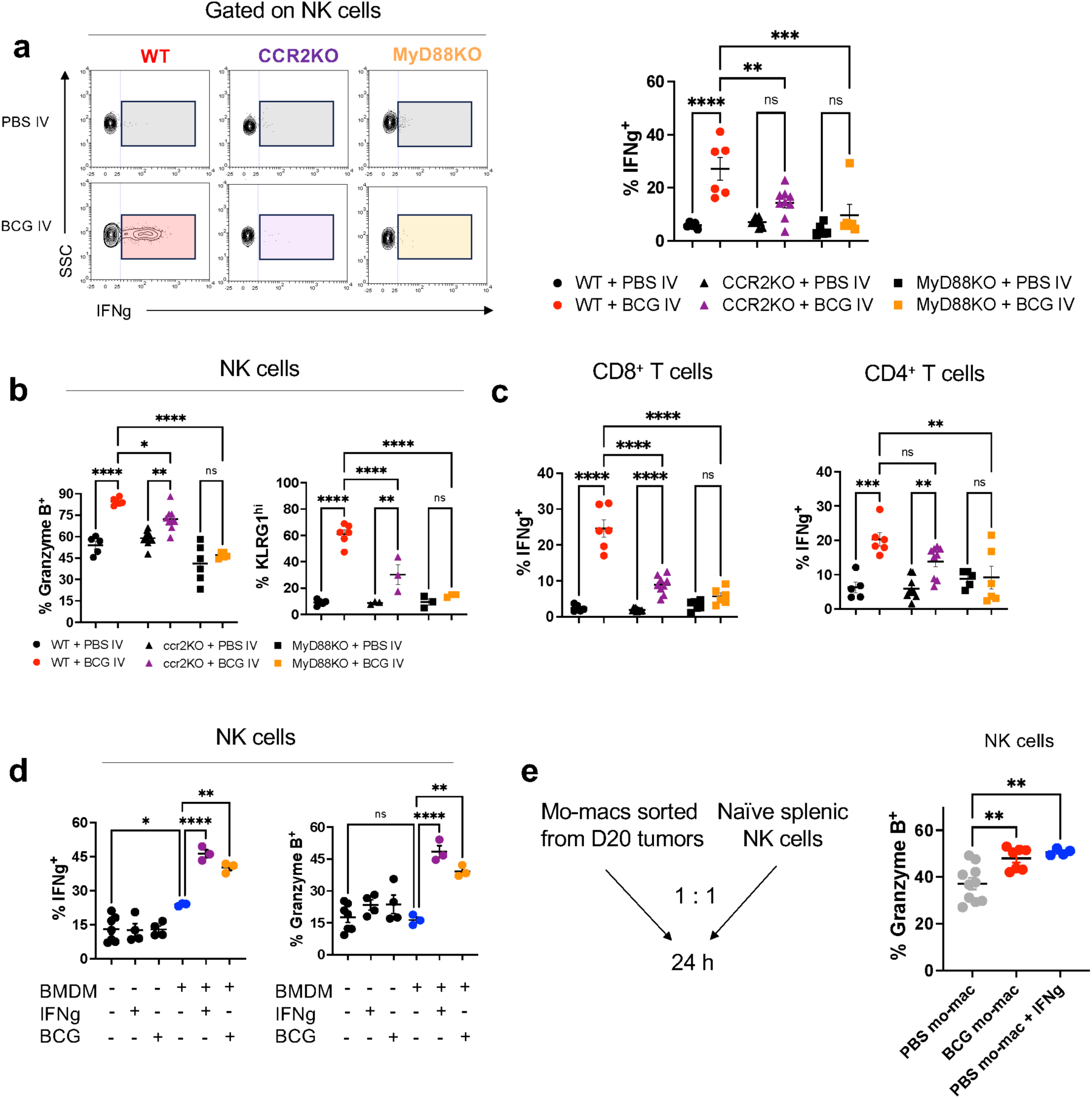
Mo-mac depletion abrogates lung NK and T cell activation by BCG. **(a)** Representative contour plots and quantification of IFNg **(a)**, Granzyme B and KLRG1 **(b)** expression by NK cells in the lungs of mice bearing B16-F10 lung tumors at day 20. **(c)** Percentage of IFNg-expressing CD8+ and CD4+ T cells in the lung. **(d)** IFNg and Granzyme B expression by *ex vivo* cultured splenic NK cells under different conditions. **(e)** Experimental setup and Granzyme B expression by ex vivo cultured splenic NK cells under different conditions. *P* values were calculated using one-way ANOVA with Bonferroni multiple-comparison test **(a,b,c,d,e)**. Significance levels are denoted as follows: ns = not significant (p ≥ 0.05), * = significant (p < 0.05), ** = highly significant (p < 0.01), *** = very highly significant (p < 0.001)

We previously showed that IV BCG treatment was not effective in IFNγ-deficient mice. Concordantly, here we found that NK cell activation by BCG was completely abrogated in the absence of IFNγ (Suppl. Fig. 6a,b). Remarkably, BCG also failed to recruit mo-macs and modulate their phenotype in these mice (Suppl. Fig. 6c,d). This result suggests that an axis involving IFNγ, recruited mo-macs and sensing of BCG via MyD88 was driving enhanced NK and T cell function in the tumor-bearing lung.

We next used an *in vitro* experimental setup to investigate whether direct interaction between macrophages and NK cells was sufficient for the activation of the latter, or additional cell types were required. First, we observed that isolated naïve NK cells were not activated by direct exposure to BCG or IFNγ, meaning that a secondary cellular mediator was needed (Fig. 5d). We then incubated bone marrow-derived macrophages (BMDMs) with naïve NK cells in the presence or absence of live BCG or IFNγ. Our data revealed that incubation of BMDMs with either IFNγ or BCG led to an increase in Granzyme B and IFNγ production by NK cells (Fig 5d).

To validate these findings in the context of tumor-bearing lungs, we incubated naïve NK cells with sorted pulmonary mo-macs (CD45^+^ CD11b^+^ F4/80^+^ CD64^+^ SiglecF^-^) from PBS- and BCG-treated tumor-bearing mice. Remarkably, mo-macs from BCG-treated group induced an increase in Granzyme B expression on the NK cells (Fig. 5e) compared to mo-macs from untreated mice, which was similar to the obtained by supplementing PBS mo-macs with IFNγ during the assay (Fig. 5e). Altogether, these *in vitro* results support our *in vivo* observations regarding the crucial role of lung mo-macs and IFNγ in triggering NK cell activation in response to IV BCG treatment.

### Reprogramming of bone marrow progenitors by BCG enhances antitumor immunity

Intravenous (IV) BCG has previously been reported to transcriptionally reprogram hematopoietic stem cells (HSCs), enhancing the generation of myeloid cells with improved capacity to control *M. tuberculosis*^17^. Notably, training of bone-marrow (BM) progenitors can also enhance antitumor responses in mouse models^18,19^. Given these findings, we hypothesized that IV BCG might also provide long-term protection against lung tumors by acting on hematopoietic progenitors in the BM. To test this hypothesis, we transplanted BM from PBS- or BCG-vaccinated mice into preconditioned recipients, which were challenged 3 months later with B16-F10 cells (Fig. 6a). As experimental controls, we first plated BM lysates to corroborate that transplanted BM did not contain viable BCG, as IV BCG can colonize the BM^17,50^ (data not shown). Additionally, using CD45.1 mice as donors to monitor transferred BM-derived immune cells, we confirmed that BM reconstitution efficiency was comparable between both groups (Suppl. Fig. 7b). Remarkably, mice reconstituted with BCG-BM exhibited significantly improved survival compared to control mice (PBS-BM) after intravenous B16-F10 challenge (Fig. 6b), as well as a reduction of lung tumor burden at day 20 (Fig. 6c). A similar effect was observed in mice transplanted with B16-F10 cells subcutaneously (Suppl. Fig. 7a), demonstrating that the antitumor effect extends beyond the lung.

**Figure 6.**
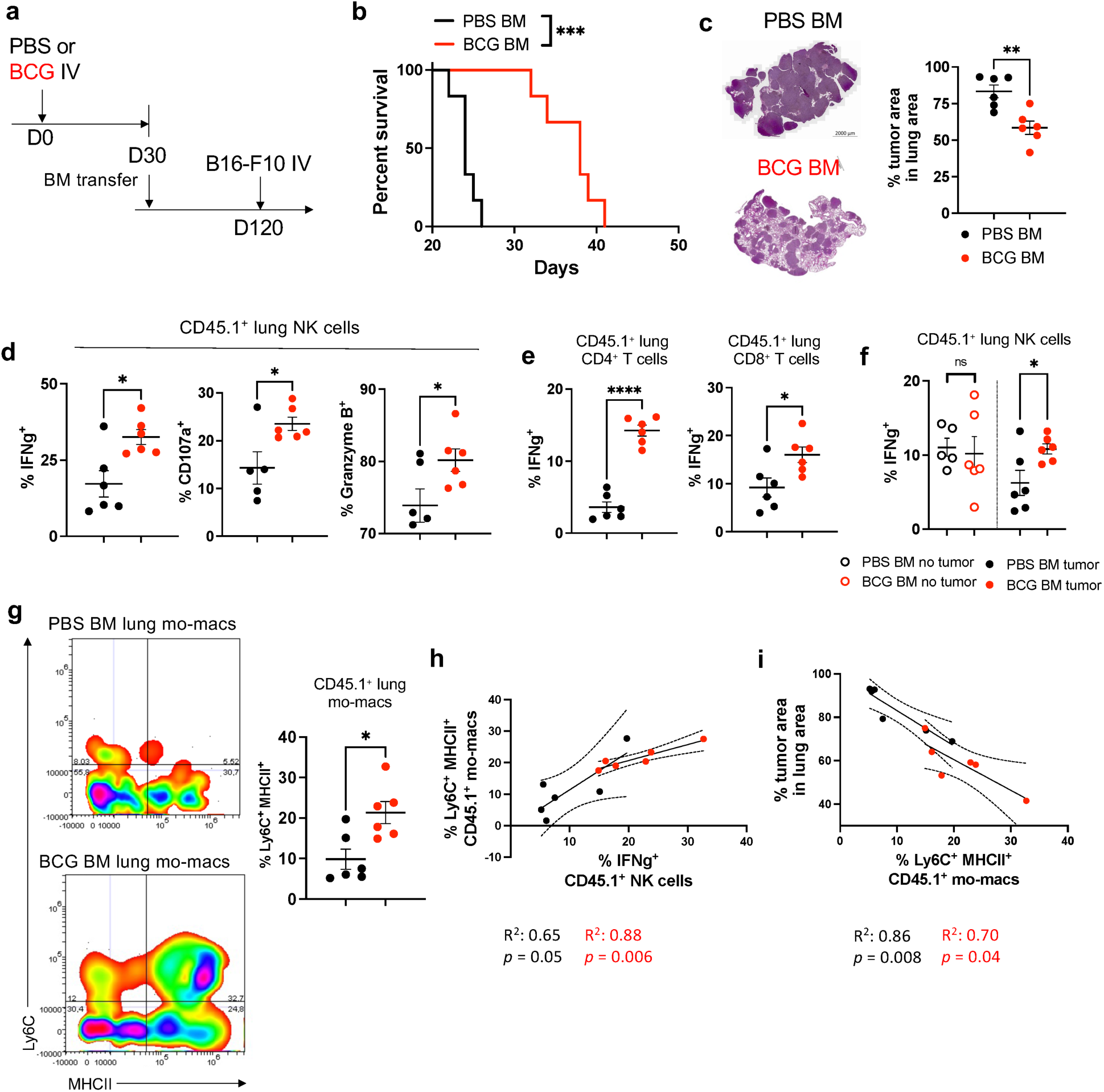
Transfer of IV BCG-conditioned bone marrow cells confers protection against lung tumor growth. **(a)** General experimental setup for bone marrow transfer experiments. **(b)** Survival of mice after B16-F10 IV inoculation. **(c)** Representative H&E images of tumor-bearing lungs at day 20 after inoculation, and quantification of the tumor area in lung cross-sections. Scale bars correspond to 1 mm in length. **(d)** Quantification of IFNg, CD107a and Granzyme B expression on CD45.1+ NK cells in the lungs of mice transplanted with PBS or BCG-BM. **(e)** IFNg expression by lung CD45.1+ CD4+ or CD8+ T cells. **(f)** IFNg expression by lung CD45.1+ NK cells. **(g)** Representative density plot and quantification of Ly6C and MHCII coexpression on CD45.1 mo-macs in the lung. **(h)** Correlation between the frequency of Ly6C+MHCII+ lung mo-macs and the frequency of IFNg+ lung NK cells. **(i)** CD86 expreesion on lung CD45.1+ cDC1s. *P* values were calculated using two-tailed unpaired Student’s t test at a 95 % CI **(c,d,e,f,g,i)** or log-rank (Mantel-Cox) test **(b)**. Significance levels are denoted as follows: ns = not significant (p ≥ 0.05), * = significant (p < 0.05), ** = highly significant (p < 0.01), *** = very highly significant (p < 0.001)

We then characterized the BM-transferred lung-infiltrating immune populations. No differences were found in the absolute numbers of different lung CD45.1 immune cell subsets (Suppl. Fig. 7c, d). Notably, CD45.1 NK and T cells in tumor-bearing lungs exhibited enhanced functionality in BCG-BM-reconstituted mice (Fig. 6d, e). Interestingly, in the absence of tumor, CD45.1^+^ NK cells displayed comparable IFNγ production between the two groups, in stark contrast to the tumor-bearing setting (Fig. 6f). This suggests that NK cell progenitors in the BM of BCG-treated mice are not intrinsically more activated, and that the observed difference is instead driven by cues within the TME.

Finally, although we did not observe differences in the total number of mo-macs between groups (Suppl. Fig. 7d), analysis of activation markers revealed a significant increase in a population of mo-macs coexpressing Ly6C and MHCII in the BCG-BM group, which was mostly absent in PBS-BM mice (Fig. 6g). Interestingly, we observed a positive correlation between NK cell activation and the presence of the Ly6C^+^ MHCII^+^ inflammatory mo-mac population in both conditions (Fig. 6h). Moreover, a higher infiltration of this mo-mac population was found correlated with reduced tumor burden (Fig. 6i) both in PBS and BCG conditions.

Altogether, these findings suggest a scenario in which IV BCG reprograms HSCs, leading—upon transfer into recipient mice—to the generation of mature mo-macs with an enhanced ability to stimulate NK and T cell antitumor responses within the lung TME.

## Discussion

Understanding the immune landscape of the tumor microenvironment requires dissecting the functional heterogeneity of its cellular components. While flow cytometry provides valuable phenotypic insights, it is inherently limited by the number of markers that can be simultaneously analyzed, making it difficult to fully resolve the complexity of immune cell states. For example, as we have demonstrated in this study, tumor-associated mo-macs exhibit a spectrum of activation states, ranging from pro-inflammatory, antitumoral phenotypes to highly immunosuppressive subsets that promote tumor progression^2,3,51^. Similarly, tumor-infiltrating T cells display functional diversity, including exhausted, dysfunctional, and effector subsets, each with distinct transcriptional signatures. By providing an unbiased, high-resolution view of immune cell states, our single-cell transcriptomic analysis uncovered key regulators of immune activation and suppression within the TME in the context of BCG treatment, offering new insights into the immune mechanisms elicited by this vaccine^52^

Our study demonstrates that IV administration of BCG exerts a strong influence in the mo-mac compartment from lung tumors and is associated with enhanced activation of NK and T cells. scRNAseq analysis revealed an enrichment of inflammatory mo-mac subsets following treatment, accompanied by a reduction or transcriptional reprogramming of immunosuppressive mo-macs. Particularly, we observed increased recruitment of classical (Mon Iigp1) and non-classical (Mon Fcgr4) monocyte populations in BCG-treated mice, which displayed a strong type II interferon-response signature with elevated expression of antigen presentation and costimulatory molecules, and immune recruiting chemokines such as *Cxcl9*. In contrast, monocyte populations in PBS-treated expressed higher levels of immunosuppressive and pro-angiogenic genes, including *Apoe, Ccl9, Fabp4, and Tgfb1*, previously linked to tumor-promoting functions^27,28,33,43^. IFN-stimulated mo-macs are known to exert potent immunostimulatory and antitumor functions^53–55^, and the ratio of Cxcl9⁺ to Spp1⁺ mo-macs has been identified as a strong prognostic marker for both overall survival and responsiveness to immune checkpoint blockade (ICB)^5^. Importantly, mature macrophages (Mac Reg) from the BCG-treated condition displayed a reduced expression of *Spp1* and *Trem2*, molecules previously involved in NK and T cell-inhibiting pathways^2,6,34^.

Interestingly, our scRNAseq data suggests that BCG has a greater influence on cellular populations that are transcriptionally closer to less differentiated monocytes than to mature macrophages. Combined with the observation that IV BCG loses its antitumor efficacy in CCR2KO mice, this finding suggests that BCG therapy primarily functions by recruiting inflammatory monocytes from the circulation, which subsequently differentiate into mature immunostimulatory mo-macs within the TME, rather than reprogramming pre-existing tumor-infiltrating macrophages, which are likely more terminally differentiated and less plastic. A previous study described an inflammatory monocyte population characterized by high expression of interferon-stimulated genes and Ly6A, capable of producing Cxcl9, Cxcl10, and Il15, and presenting tumor antigens via cross-dressing^37^. In our dataset, we also identified *Ly6a* as a defining feature of the Mon Iigp1 and Mon Fcgr4 populations enriched in BCG-treated mice. However, in contrast to the aforementioned study, this *Ly6a* population in our model was associated with type II IFN responses rather than type I IFN signaling.

Ligand-receptor analysis revealed a central role for pro-inflammatory mo-mac populations from IV BCG-treated mice in engaging NK cells and T lymphocytes. Notably, T cells are required for the antitumor effects of both intravesical BCG in bladder cancer^56^ and IV BCG in lung tumors^16^, as well as for tuberculosis protection in macaques^14^. But, how are NK and T cells initially activated in the lung tumor microenvironment following IV BCG? Our results show that after IV administration of BCG-GFP, bacilli are predominantly found within the lung mo-mac compartment. We propose that early recognition of BCG by mo-macs triggers a primary MyD88-dependent innate inflammatory response, initiating the recruitment of additional mo-macs to the lung via CCL2-CCR2 signaling. Upon encountering BCG-infected macrophages, NK and T cells begin producing IFN-γ and other cytokines, initiating a second, delayed wave of inflammation. This response further recruits mo-macs and modulates their phenotype through IFNγ signaling. IFNγ-licensed mo-macs would then produce inflammatory mediators such as IL-15, IL-27, CXCL9/10 and CCL5, which further enhance and sustain NK and T cell activation and recruitment in a feed-forward loop. Consistent with this model, the absence of IFN-γ, MyD88 or CCR2 completely abolishes the IV BCG-induced activation of both mo-macs and NK cells, and addition of BCG or IFNγ to BMDM-NK cell co-cultures induces naïve NK cell activation. This MyD88-dependent biphasic immune activation pattern was previously observed in the context of IV BCG-mediated heterologous protection against influenza infection^57^, although the specific contribution of mo-macs was not assessed.

Further supporting the point that IV BCG therapy mainly functions by recruiting inflammatory monocytes from the circulation, we show that mice transplanted with BCG-conditioned bone marrow—an experimental setting in which the immediate inflammatory effects derived from the direct presence of BCG are absent—still mounted enhanced antitumor responses. The observed effect in this scenario may also be driven by the recruitment of BM-derived immunostimulatory mo-macs to the tumor, which in turn promote NK and T cell activity. Recent studies have shown that training of BM progenitors by intravesical or IV BCG can boost antitumor immunity in the bladder and subcutaneously-implanted tumors^58,59^. However, those effects were primarily attributed to neutrophils rather than mo-macs. In our dataset, we also observed a type II interferon signature in lung neutrophils, although we did not directly assess their contribution to the antitumor response. Interestingly, the mentioned studies reported an increased MHC class II expression in monocytes from BCG-experienced mice, although their functional contribution to the antitumor effect was not evaluated. Taken together, these findings in conjunction with ours suggest that training of BM progenitors can give rise to both neutrophil and mo-mac populations with enhanced anti-tumor properties. While trained neutrophils contribute to antitumor immunity via ROS production and direct tumor cell killing, trained mo-macs seem to work mainly by favoring the activation of other immune cell subsets such as NK or T cells. The capacity of IV BCG to remodel BM progenitors contrasts with the intrinsic ability of tumors to modulate the BM compartment via cytokines such as IL-6 or IL-4^60,61^, leading to the production of immunosuppressive myeloid cells. It is tempting to speculate that the ability of BCG to alter HSCs via T cell and NK cell-derived IFNγ^17^ can counter the IL-4 and IL-6-mediated effects exerted by the tumor, and thus abrogate the generation of myeloid cells with an immune-suppressive phenotype. Importantly, an effect of BCG on HSCs has also been observed in humans^24^.

Traditional checkpoint blockade therapies primarily target exhausted T cells rather than modulating the myeloid compartment^62,63^. Checkpoint blockade shows limited efficacy in cold tumors infiltrated with poorly activated T cells, as is the case for B16-F10 lung metastases^64^. Unlike PD-1 or CTLA-4 inhibitors, IV BCG, reminiscent of other innate immune system-targeting therapies such as TLR agonists or CD40-targeted antibodies^65^, appears to act upstream by remodeling the immune landscape of tumors, creating a more favorable environment for lymphocyte activation. However, its unique ability to persist in the host and to modulate BM progenitors, leading to the generation of myeloid cells with a higher immunostimulatory profile, might confer durable protection against tumor recurrence in the long term. Our work underscores the therapeutic potential of cancer immunotherapies that engage both the myeloid and lymphoid compartments to achieve effective and sustained antitumor responses.

## Methods

### Mouse strains

Female mice between the ages of 8 and 12 weeks were used. C57BL/6JR mice were purchased to Janvier Biolabs. Mouse strains deficient for interferon gamma, CCR2 and MyD88 on C57BL/6JR background were bred in the facilities of Centro de Investigaciones Biomédicas de Aragon. CD45.1 mice (B6.Cg-Ptprca-Pepcb/Rj) were purchased to Janvier Biolabs. Mouse experimentation and breeding were performed in a SPF-facility at 20-24 °C, 50-70 % humidity and a light-dark cycle of 12 h.

### Tumor cell lines and tumor outgrowth studies

B16.F10 tumor cells were cultured with complete DMEM, containing 10% inactivated Fetal Bovine Serum (FBS, Gibco), Glutamax (Gibco) and penicillin/streptomycin (Gibco) and were always used with less than 8 passages from thawing. For lung tumor induction, 1 × 10^5^ B16-F10 cells were injected intravenously in PBS. For flow cytometry and scRNAseq analyses, mice were sacrificed at day 20 post tumor inoculation. For survival experiments, mice were sacrificed based on a clinical score including evaluation of weight loss (a maximum of 20% of initial weight loss was allowed), general appearance and activity, and the frequency and quality of breathing, which was performed every 2 days for the duration of the experiment.

For subcutaneous tumors, 5 x10^5^ B16-F10 cells resuspended in PBS were injected in the flank. Size of subcutaneous tumors was measured with a digital caliper two-three times per week and determined by using the following formula: [(tumor width)^2^× (tumor length)]/2. Mice were sacrificed by inhaled anesthesia followed by cervical dislocation either when tumor volume exceeded 1 cm^3^, when tumor diameter exceeded 10 cm in any direction, or when tumors became ulcerated.

### Bacterial strains

The BCG Pasteur 1173P2 (a kind gift from Dr. Roland Brosch, Institute Pasteur, France) used in this study was grown at 37 °C in Middlebrook 7H9 broth (BD Difco) supplemented with 0.05% Tween 80 (Sigma) and 10% Middlebrook albumin dextrose catalase enrichment (ADC; BD Biosciences). A solution containing 10^6^ CFUs of BCG Pasteur diluted in PBS was inoculated intravenously.

### Clodronate liposome treatments

For circulating monocyte depletion, 200 μl of either control or clodronate liposomes (LIPOSOMA) were injected intravenously every 3 days until the end of the experiment.

### Preparation of single-cell suspensions

Tumor-bearing lungs were aseptically removed and homogenized in DMEM containing deoxyribonuclease I (DNase I, 40 U ml^−1^; AppliChem), collagenase D (2 mg ml^−1^; Roche) and Liberase TM (0,2 U ml^−1^) using a GentleMacs dissociator (Miltenyi Biotec) according to manufacturer’s instructions. Lungs were incubated at 37 °C for 30 min and further homogenized with the GentleMacs dissociator. The homogenates were filtered through a 70 µm cell strainer (MACS SmartStainers, Miltenyi Biotec). Erythrocytes were lysed with RBC Lysing Buffer for 1 min and single cells were resuspended in PBS with 2% FBS and 2 mM EDTA and stained for surface and intracellular markers.

### Re-stimulation and intracellular cytokine staining

For T cells, lung single cell suspensions were stimulated with 2 µg ml^−1^ of plate bound αCD3 (Miltenyi, clone 145-2C11) and 5 µg ml^−1^ of soluble αCD28 (BD, clone 37.51) for 4 h at 37 °C in the presence of Brefeldin A (eBioscience) in complete RPMI medium containing 50 µM β-mercaptoethanol (Gibco). For ex vivo NK cell stimulation, single cell suspensions were cultured with 20 µg ml^−1^ of plate bound αNK1.1 (clone PK136) in the presence of FITC-conjugated anti-CD107a antibody (BD, clone 1D4B) and Brefeldin A for 4 h at 37 °C.

### Flow cytometry

Single cells were incubated with mouse Fc receptor blocking reagent (Miltenyi) for 20 min at 4 °C, washed and stained with fluorochrome-conjugated antibodies (at a 1:200 dilution, Supplementary Table 1 and Supplementary Table 4) for 20 min at 4 °C. Cells were fixed in 4 % (v/v) histoloy grade Formalin and acquired in a Gallios flow cytometer (Beckman Coulter) or an Aurora Cytek 5 L spectral flow cytometer. For intracellular staining, cells were first fixed in 2 % (v/v) histoloy grade Formalin and then permeabilized with the eBioscience Foxp3 / Transcription Factor Staining Buffer Set according to manufacturer instructions. Gating strategies to define immune cell populations are described in Supplementary Figs. 8 and 9.

### TAM sorting

Single cell suspensions of lungs were obtained and stained as described. TAMs were sorted as CD45^+^ SiglecF^-^ CD11b^+^ F4/80^+^ in a SH800S Cell Sorter (Sony). Sorted cells were centrifuged and resuspended in RPMI containing 10% inactivated Fetal Bovine Serum (FBS, Gibco), Glutamax (Gibco) and penicillin/streptomycin (Gibco). Sorted TAMs were left to adhere overnight.

### BMDM generation

Bone-marrow derived macrophages (BMDMs) were generated from the tibias and femurs of female mice. Bone-marrow single cell suspensions were cultured for 7 days in complete DMEM supplemented with 20 % of supernatant from L929 fibroblasts as a source of M-CSF.

### Macrophage-NK cell cocultures

Untouched NK cells were isolated from the spleens of naïve mice by magnetic separation with the EasySep mouse NK cell isolation kit. 1x10^5^ NK cells were cocultured with 1x10^5^ TAMs or BMDMs in complete RPMI in 96 well plates for 24h in the presence or absence or live BCG Pasteur (at a MOI of 1 bacteria per macrophage), IFNγ (10 ng ml^-1^, Peprotech) or IL-15 (10 ng ml^-1^). Then, NK cells were transferred to another 96 well plate and stimulated with plate-bound antiNK1.1 (20 μg ml^-1^) for 4h in the presence of brefeldin A in complete RPMI, and stained for extracellular and intracellular markers as described.

### Bone-Marrow cell isolation and transfer

CD45.1 mice were treated with IV BCG or control IV PBS as has been previously described. 30 days after treatment, femurs and tibias were extracted and the bone marrow cells isolated as previously described. 2,5x10^6^ cells were transferred into recipient CD45.2 mice which had previously been treated with 20 mg/kg of 1,4-Butanediol dimethanesulfonate (busulfan) intraperitoneally for 4 consecutive days to deplete their bone marrow^66^. Bone-marrow reconstitution was allowed for 3 months and then mice were used for subsequent experiments.

### Histology

Tumor quantification was performed in formalin fixed paraffin-embedded lung tissue sections (4 µm) mounted in slides which were digitalized with a Zeiss AxioScan slide scanner and analyzed using ImageJ software.

### scRNAseq samples and library preparation

Lungs from five mice per experimental group (BCG-treated or PBS-treated) were perfused with cold PBS and single cell suspensions prepared as described above. Single cell suspensions were stained with anti-CD45 antibody for 20 minutes on ice, and CD45⁺ cells were separated in a SH800S Cell Sorter (Sony). The isolated cells were processed on the 10x Genomics Chromium Controller to generate GEM bead emulsions with the Single Cell 3′ Library & Gel Bead Kit v3.1 (10x Genomics), making calculations for 10.000 cells/library. cDNA synthesis, amplification, and subsequent library preparation were carried out according to the manufacturer’s protocols. For sequencing, both libraries were pooled and sequenced on a NovaSeqX platform (Illumina), targeting an average read depth of 40.000 reads per cell. 1% of PhiX was added to the lane to ensure diversity.

### Data processing and filtering

Raw data (fastqs) were processed using Cell Ranger pipeline (v7.2.0, Transcriptome; mm10-2020-A). Filtered gene-expression matrices were subjected to processing by Seurat R Package (v 5.1.0). Quality control measures were implemented to remove debris, damaged cells and doublets (min_genes_per_cell = 300; max_genes_per_cell = 5500; max_%_mitochondrial_reads = 20), and batch effect was corrected using SCTransform approach^67^. We generated a BCG single-cell library containing 16,507cells and a PBS single-cell library containing 12,019 cells, with a median of over 2,000 expressed genes per cell in both conditions.

### scRNAseq clustering and analysis

Principal component analysis (PCA) was performed and UMAP dimensionality reduction was subsequently applied based on the first 30 dimensions. Neighbor identification was performed using the first 30 principal components, followed by clustering using the Louvain algorithm (resolution = 0.5) in Seurat. Differential expression analysis identified the top-ranked genes (hub genes) that were upregulated in each individual cluster relative to the combination of all other cells, as determined by FindAllMarkers in Seurat. Annotation of each unbiased population was achieved through manual inspection of the top-ranked genes of each cluster. For naïve T cells, we used FindSubCluster (resolution = 0.1, Louvain algorithm) from Seurat to differentiate between CD4 and CD8 subsets. Gene scores were computed using the AddModuleScore function from Seurat with curated gene lists provided. Statistical analyses were then performed using Wilcoxon tests, with p-value adjustments applied using the Bonferroni method.

### Statistical analysis

Comparisons between clusters of interest were conducted using the FindMarkers function from Seurat (Wilcoxon tests with Bonferroni post-testing) with the options logfc.threshold = 0, min.pct = 0, and min.diff.pct = - Inf to retain all available data. Filtered results were subsequently generated using in-house scripts, applying a minimum adjusted p-value threshold of 0.05 and a minimum log2 fold change of 0.5. The results were visualized using the EnhancedVolcano R package, displaying the top 50 genes ranked by adjusted p-value. For studying shared and exclusive genes between populations, we used nvennR as authors recommended for visualization and exploration of results^68^.

### Gene ontology and gene set enrichment analysis

ClueGO plugin (v2.5.10) was utilized in Cytoscape software (v3.10.2) with default settings and the GO-ImmuneSystemProcess-EBI-UniProt-GOA-ACAP-ARAP-25.05.2022 database. Heatmaps were generated using ggplot2, based on adjusted p-values (-log10 transformation), with hierarchical clustering performed using Euclidean distances and the Ward.D2 method to group GO terms by similarity. GSEA was performed using the fgsea R package (v1.26.0)^69^ on ranked gene lists derived from differential expression data. Reactome pathways from the msigdb_v2024.1.Mm_GMTs collection were used as the reference gene sets. Significant pathways (adjusted p-value ≤ 0.05) were visualized using ggplot2.

### Cell-Cell communication network, Trajectory and Pseudotime analysis

Cell-Cell interactions were inferred using CellChat (v2.1.2) following official tutorials with the database CellChatDB.mouse containing “Secreted Signaling”, “ECM-Receptor” and “Cell-Cell Contact” categories^38^. For trajectory and pseudotime analysis we employed Monocle3 (v1.3.7) as authors recommended^70^. We imported our Seurat object to Monocle3 using SeuratWrappers package in R (v0.3.5). Additionally, transcriptional dynamics were analyzed using RNA velocity with scVelo (v0.3.3)^71^, in combination with Scanpy (v1.11.0) and Anndata (v0.11.3)^72^. Loom files were generated using velocyto (v0.17.17) from aligned BAM files and genome annotations (mm10). AnnData objects were filtered, normalized, and processed following current scVelo guidelines, including moment computation, stochastic velocity estimation, and velocity graph construction. Trajectories were visualized by projecting velocity vectors onto the precomputed UMAP embedding.

## Author approval

All the authors have revised the manuscript and approve this submission

## Acknowledgments

This work was supported by MCIN/AEI/10.13039/501100011033 [grants number RTI2018-097625-B-I00 and PID2022-138624OB-I00], Gobierno de Aragón [grant number LMP50_21] and Asociación Española Contra el Cáncer (AECC) [grant number IDEAS211042AGUI]. This work has been co-financed by the Spanish Ministry of Science and Innovation with funds from the European Union NextGenerationEU, from the Recovery, Transformation and Resilience Plan (PRTR-C17.I1) and from the Autonomous Community of Aragón within the framework of the Biotechnology Plan Applied to Health. NA was the principal investigator of all these grants. This research was supported by CIBER-Consorcio Centro de Investigación Biomédica en Red-(Groups CB06/06/0020), Instituto de Salud Carlos III, Ministerio de Ciencia e Innovación and Unión Europea-European Regional Development Fund. CG has a pre-doctoral fellowship from Gobierno de Aragón. The funders had no role in study design, data collection and analysis, decision to publish or preparation of the manuscript. Authors acknowledge the Scientific and Technical Services from Instituto Aragonés de Ciencias de la Salud (IACS) and Universidad de Zaragoza.

## Supplementary Figures

**Supplementary Figure 1.**
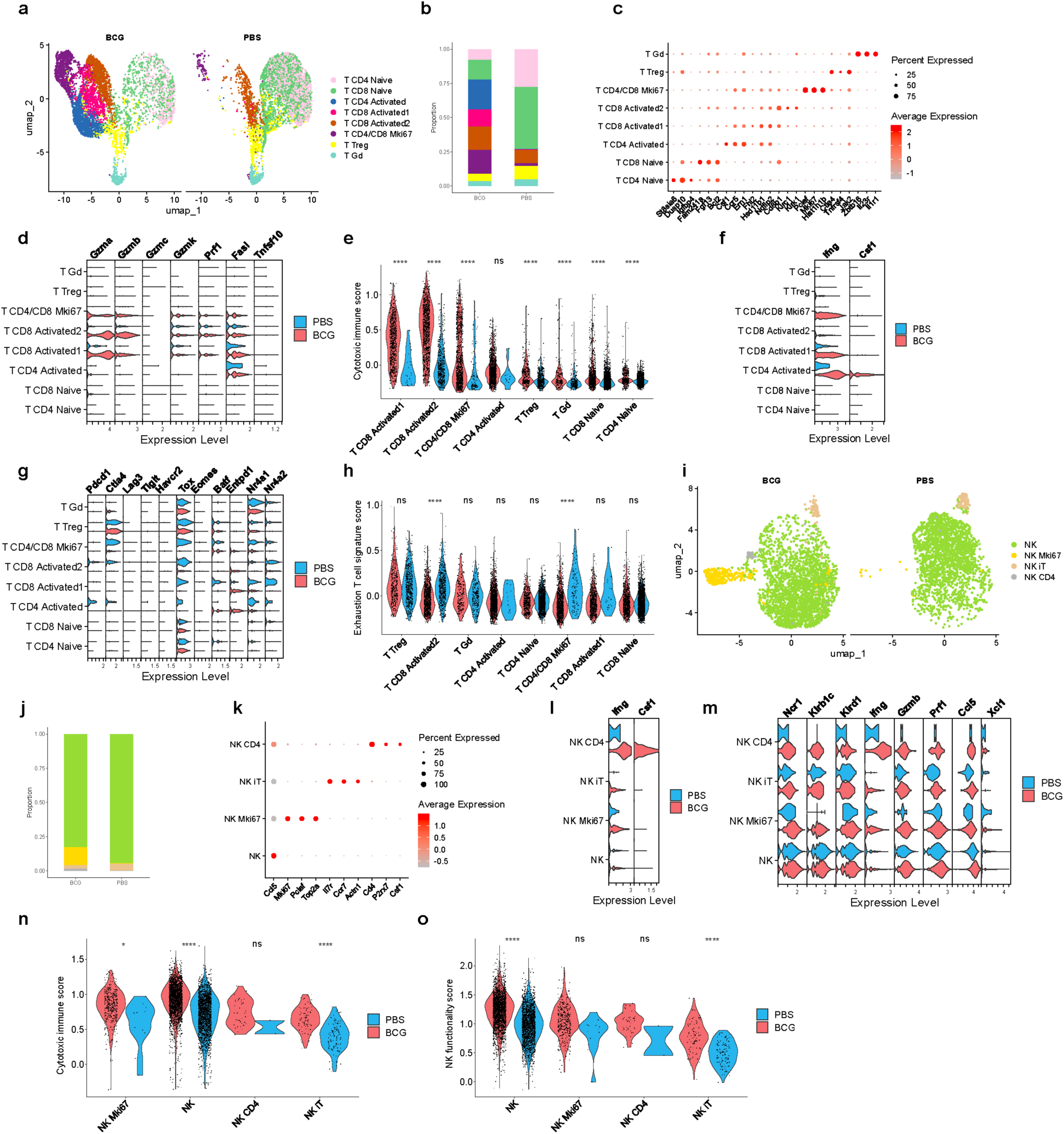
Subclustering analysis of T and NK cell subpopulations in tumors from B16F10-bearing mice. (**a**) UMAP plot, (**b**) proportions, (**c**) hub genes, (**d**) functional markers, (**e**) cytotoxic immune score, (**f**) expression level of Ifng and Csf1 transcripts, (**g**) expression level of curated genes related to exhaustion, and (h) exhaustion T cell signature score in T cells. (**i**) UMAP plot, (**j**) proportions, (**k**) hub genes, (**l**) expression level of Ifng and Csf1 transcripts, (**m**) functional markers, (**n**) cytotoxic immune score, and (**o**) NK functionality score in natural killer cells. Statistical analysis were performed using Wilcoxon tests, with p-value adjustment applied using the Bonferroni method. Significance levels are denoted as follows: ns = not significant (p ≥ 0.05), * = significant (p < 0.05), ** = highly significant (p < 0.01), *** = very highly significant (p < 0.001), and **** = extremely significant (p < 0.0001).

**Supplementary Figure 2.**
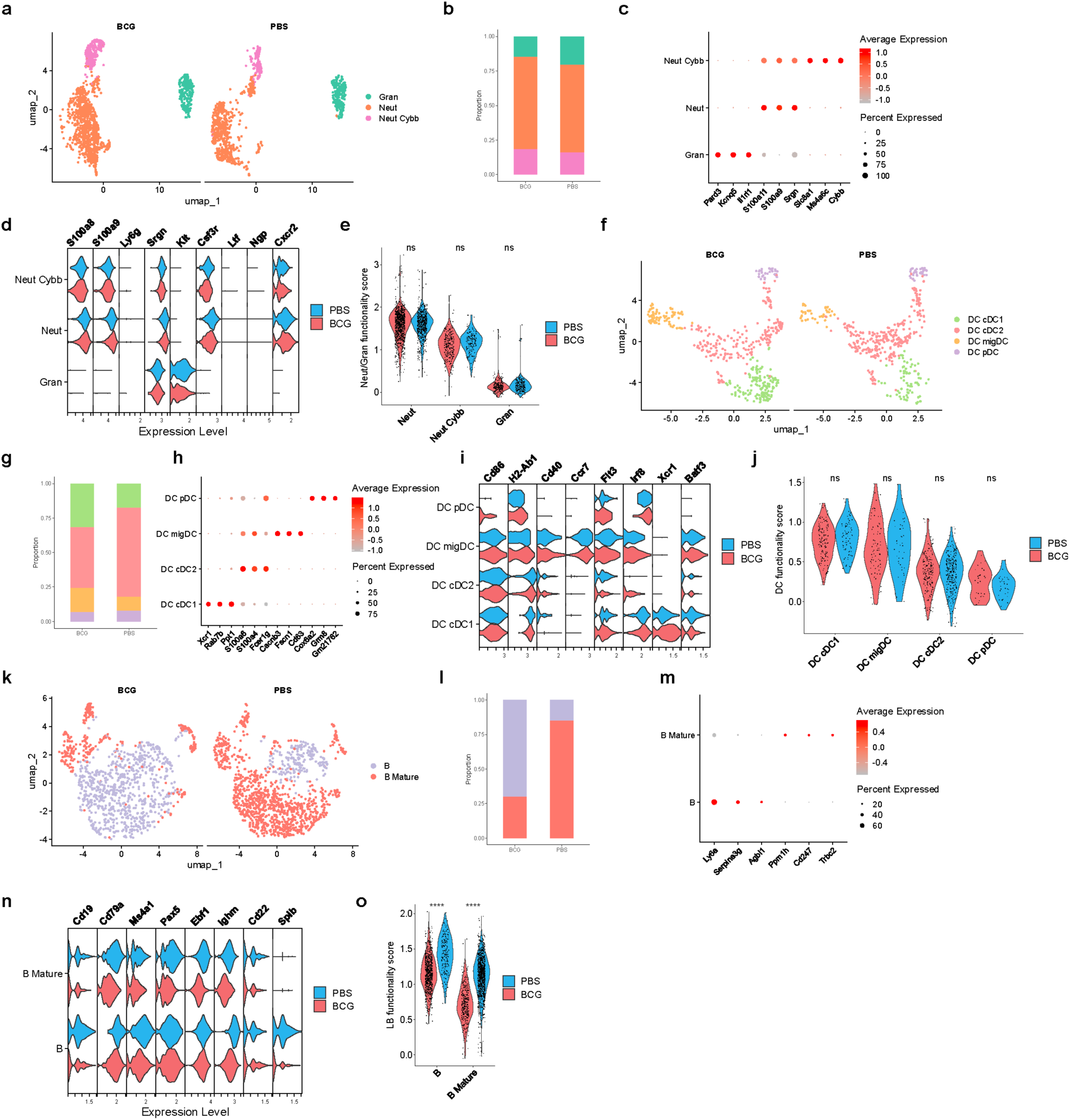
Subclustering analysis of neutrophils, granulocytes, dendritic cells and lymphocytes. **B.** (**a**) UMAP plot, (**b**) proportions, (**c**) hub genes, (**d**) functional markers and (**e**) functionality score in neutrophils and granulocytes. (**f**) UMAP plot, (**g**) proportions, (**h**) hub genes, (**i**) functional markers and (**j**) functionality score in dendritic cells. (**k**) UMAP plot, (**l**) proportions, (**m**) hub genes, (**n**) functional markers and (**o**) functionality score in dendritic cells. Statistical analysis was performed using Wilcoxon tests, with p-value adjustment applied using the Bonferroni method. Significance levels are denoted as follows: ns = not significant (p ≥ 0.05), * = significant (p < 0.05), ** = highly significant (p < 0.01), *** = very highly significant (p < 0.001), and **** = extremely significant (p < 0.0001).

**Supplementary Figure 3.**
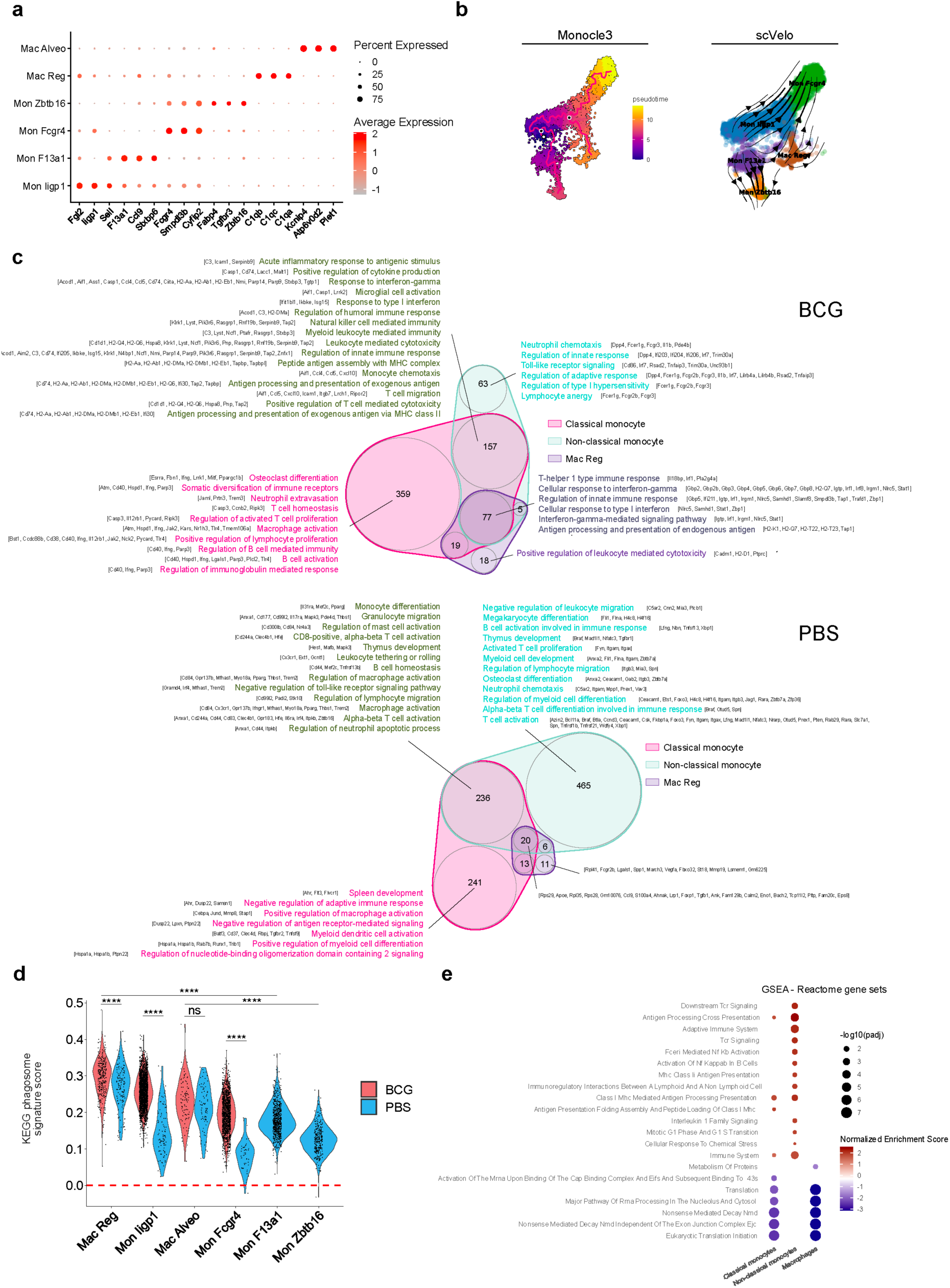
Transcriptional characterization and dynamics of mo-macs subpopulations in B16F10 tumors upon BCG treatment from B16F10-bearing mice. (**a**) Hub genes defining mo-macs populations across conditions. (**b**) Pseudotime and trajectory analysis of mo-macs subsets using Monocle3 and scVelo. (**c**) Venn diagram showing shared and unique DEGs among mo-macs subsets. GO analysis of unique and shared DEGs highlights enriched functional categories. (**d**) Violin plot showing KEGG phagosome (mmu04145) signature scores across mo-macs subsets.

**Supplementary Figure 4.**
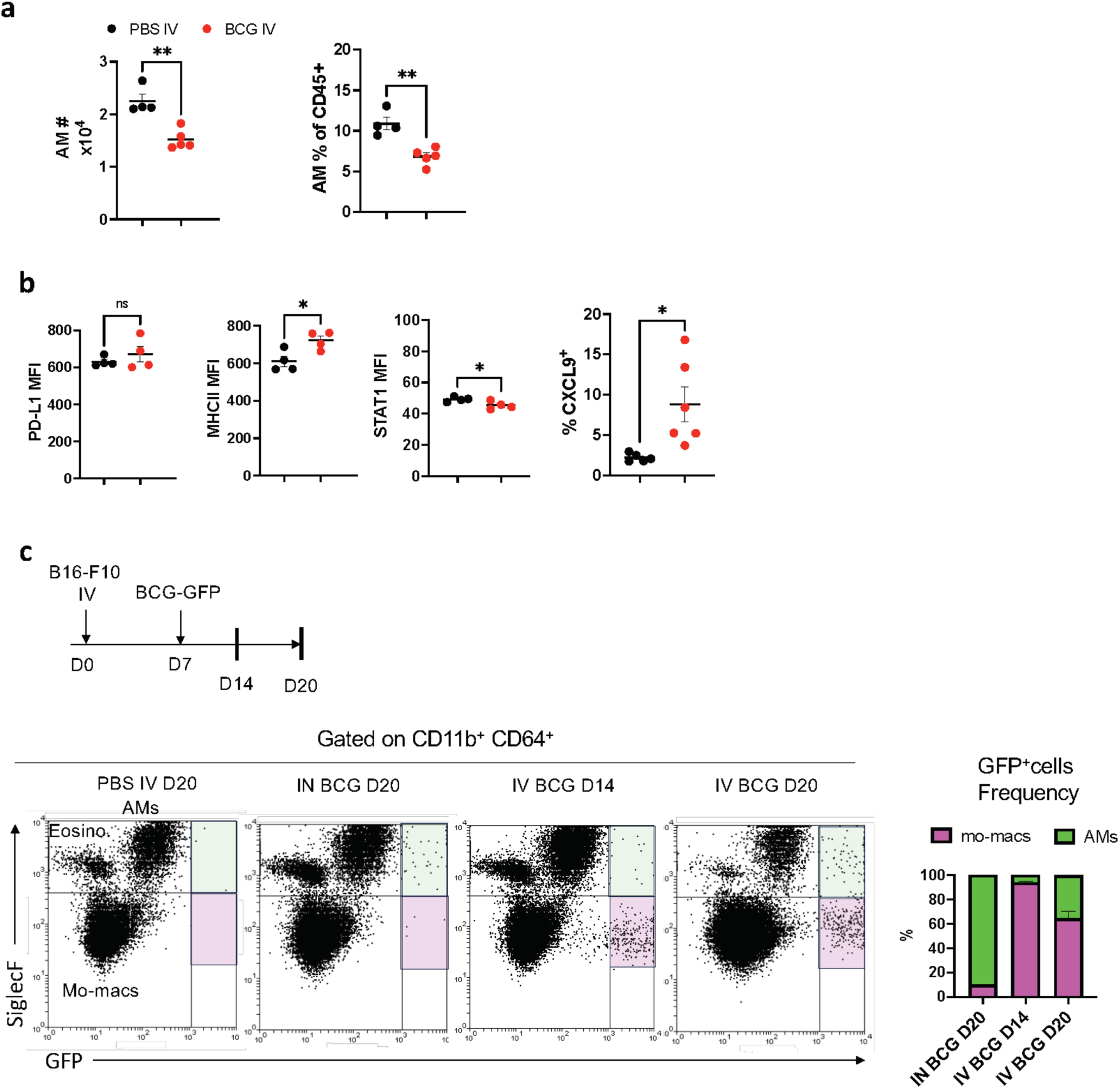
Effect of IV BCG on AMs and uptake of BCG by distinct lung macrophage populations. **(a)** Absolute number and frequency of alveolar macrophages in the lungs of mice bearing B16-F10 tumor at day 20, analyzed by flow cytometry in single cell suspensions**. (b)** Expression of surface and intracellular markers on AMs. **(c)** Experimental setting for mice treatment with GFP-expressing BCG, representative plots of GFP expression on lung CD11b+ CD64+ myeloid cells and quantification. *P* values were calculated using two-tailed unpaired Student’s t test at a 95 % CI **(a,b)**. Significance levels are denoted as follows: ns = not significant (p ≥ 0.05), * = significant (p < 0.05), ** = highly significant (p < 0.01), *** = very highly significant (p < 0.001).

**Supplementary Figure 5.**
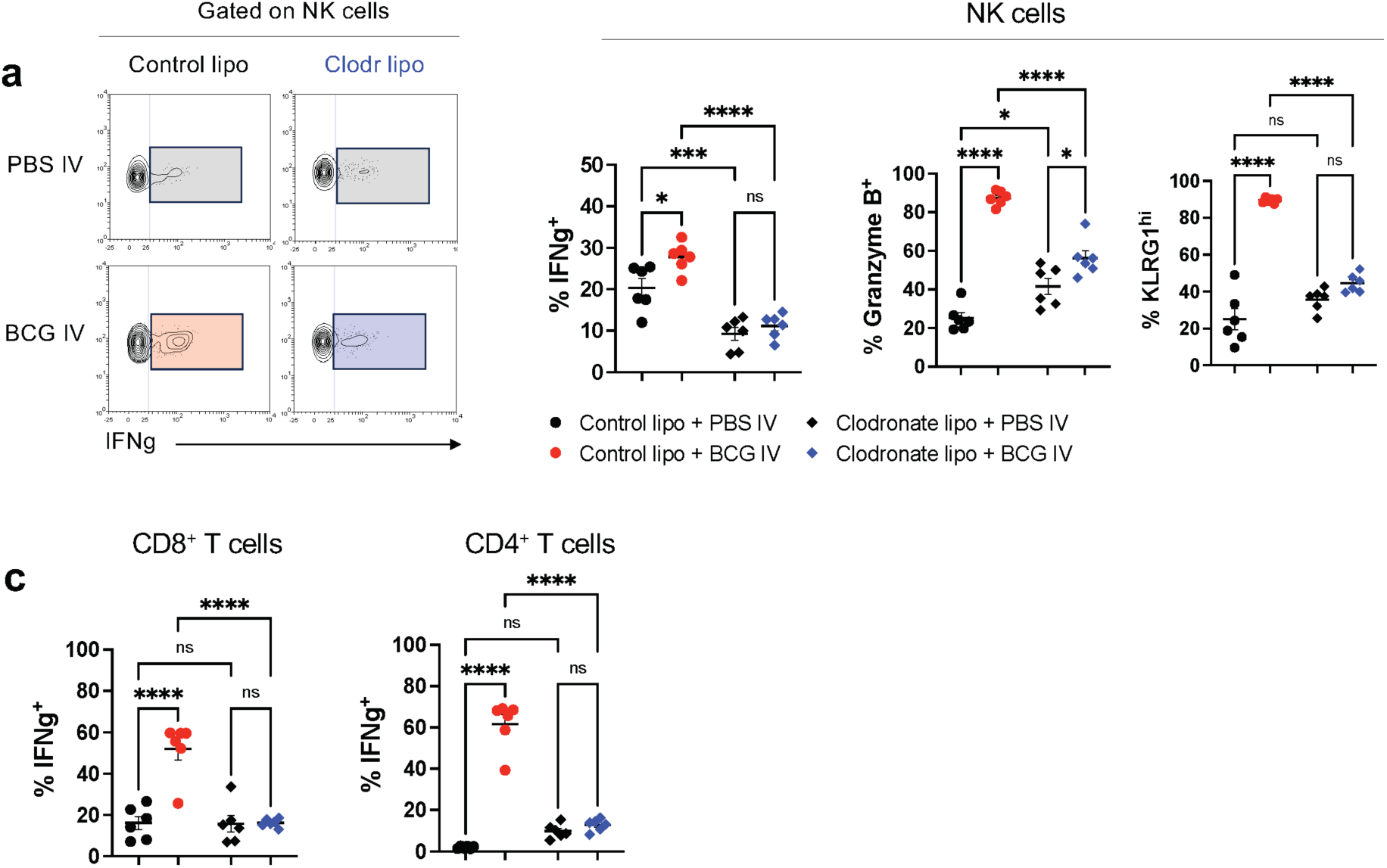
Clodronate liposome treatment abrogates NK and T cell activation by IV BCG. **(a)** Representative dot plots and quantification of IFNg, Granzyme B and KLRG1-expressing NK cells **(a)** and IFNg in T cells **(b)**, from mice treated as in Figure 4f. *P* values were calculated using one-way ANOVA with Bonferroni multiple-comparison test **(a,b)**. Significance levels are denoted as follows: ns = not significant (p ≥ 0.05), * = significant (p < 0.05), ** = highly significant (p < 0.01), *** = very highly significant (p < 0.001).

**Supplementary Figure 6.**
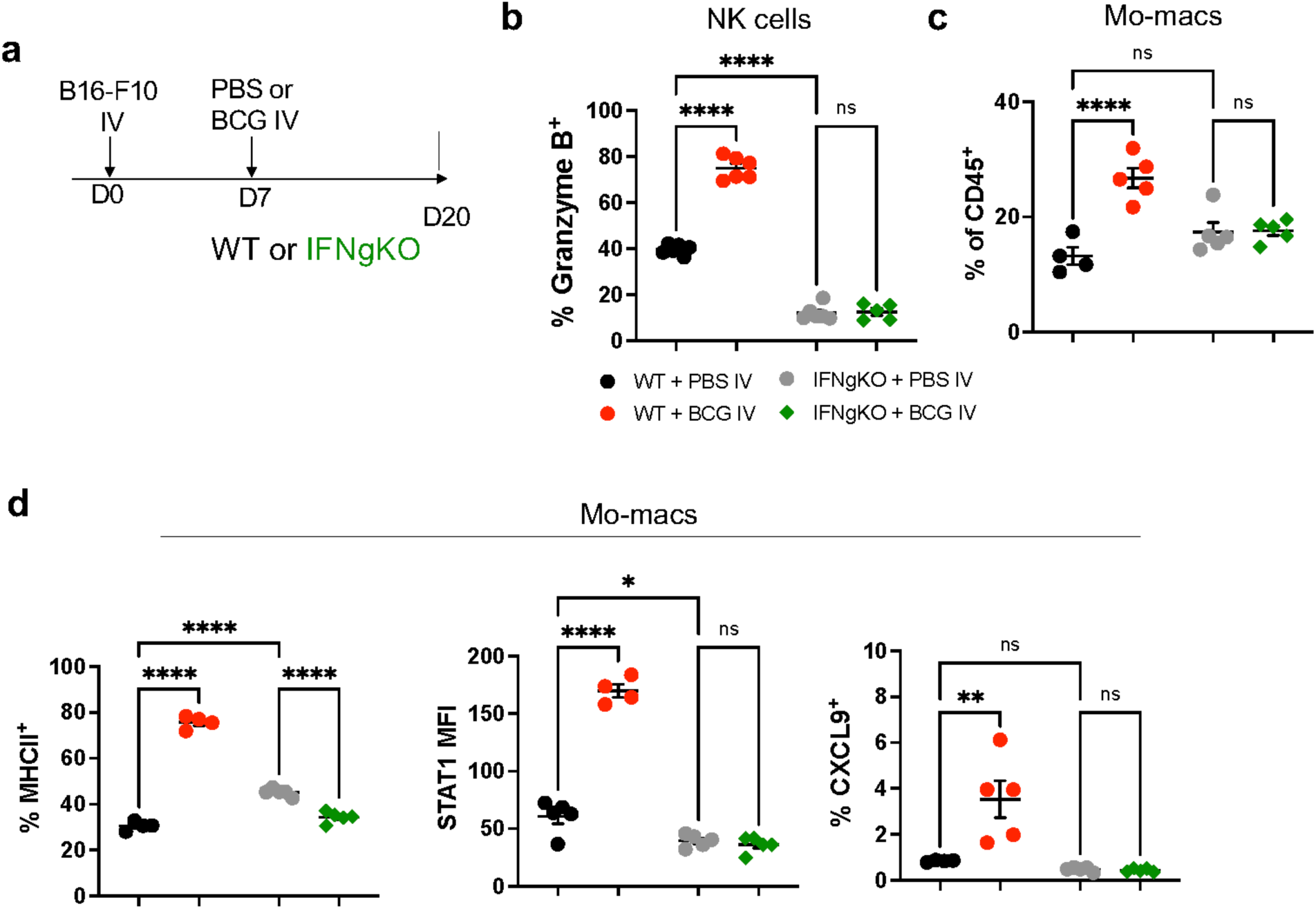
Absence of NK cell and mo-mac activation in IFNgKO mice. **(a)** Experimental setup. **(b)** Quantification of Granzyme B expression by lung NK cells. **(c)** Frequency of mo-macs in the lungs. **(d)** Extracellular and intracellular marker expression on lung mo-macs. *P* values were calculated using one-way ANOVA with Bonferroni multiple-comparison test **(b,c,d)**. Significance levels are denoted as follows: ns = not significant (p ≥ 0.05), * = significant (p < 0.05), ** = highly significant (p < 0.01), *** = very highly significant (p < 0.001).

**Supplementary Figure 7.**
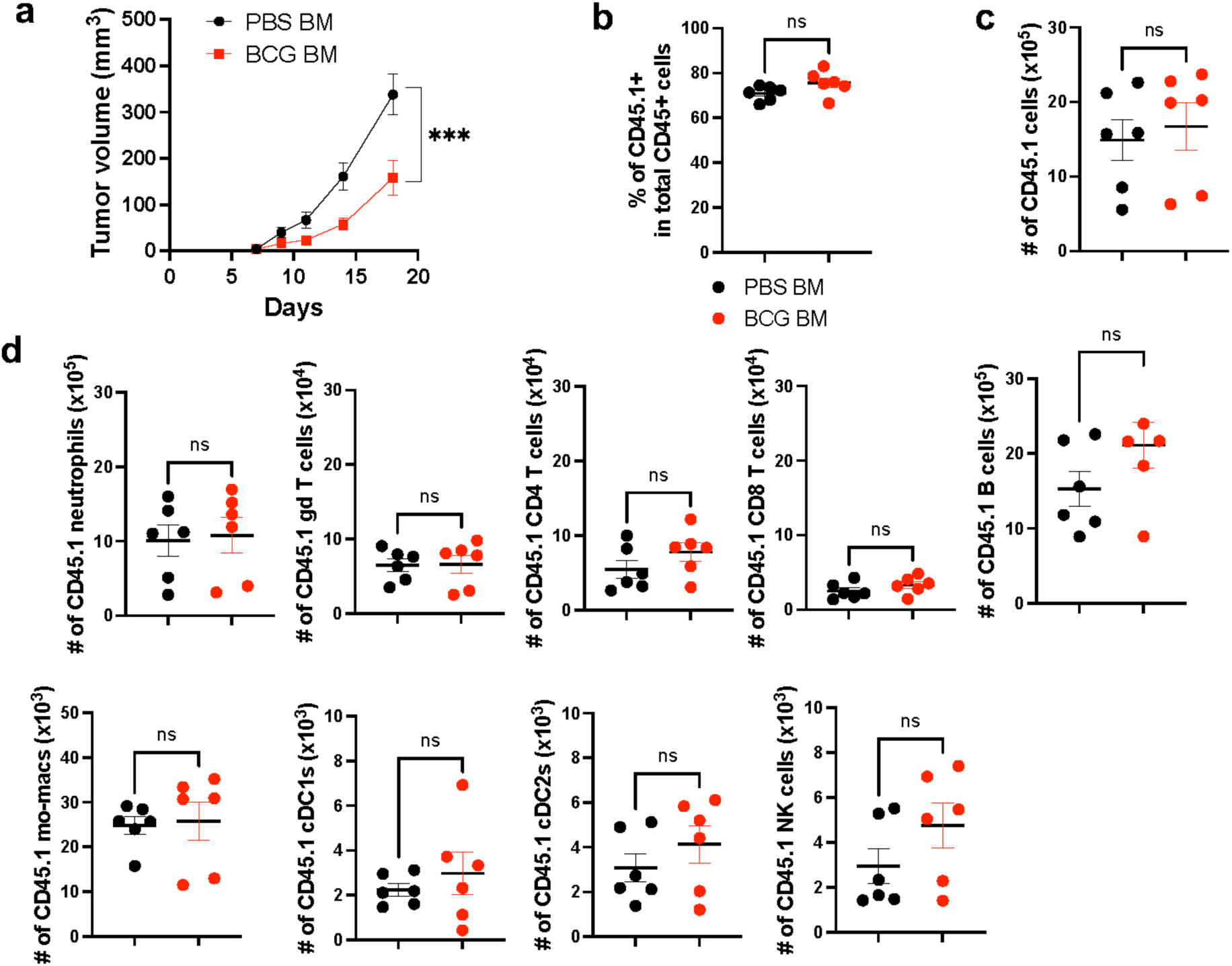
Related to figure 6 – Transfer of IV BCG-trained bone marrow cells confers protection against lung tumor growth. **(a)** Frequency of CD45.1+ cells among total CD45 cells in the lungs of mice at day 20 post tumor implantation**. (b)** Tumor growth curves of mice subcutaneously transplanted with B16-F10 cells. **(c,d)** Absolute numbers of distinct CD45.1+ immune cell subsets in the lungs of mice at day 20 post tumor implantation.. *P* values were calculated using two-way repeated measures ANOVA **(a)** or two-tailed unpaired Student’s t test at a 95 % CI **(b,c,d,e).** Significance levels are denoted as follows: ns = not significant (p ≥ 0.05), * = significant (p < 0.05), ** = highly significant (p < 0.01), *** = very highly significant (p < 0.001).

**Supplementary Figure 8.**
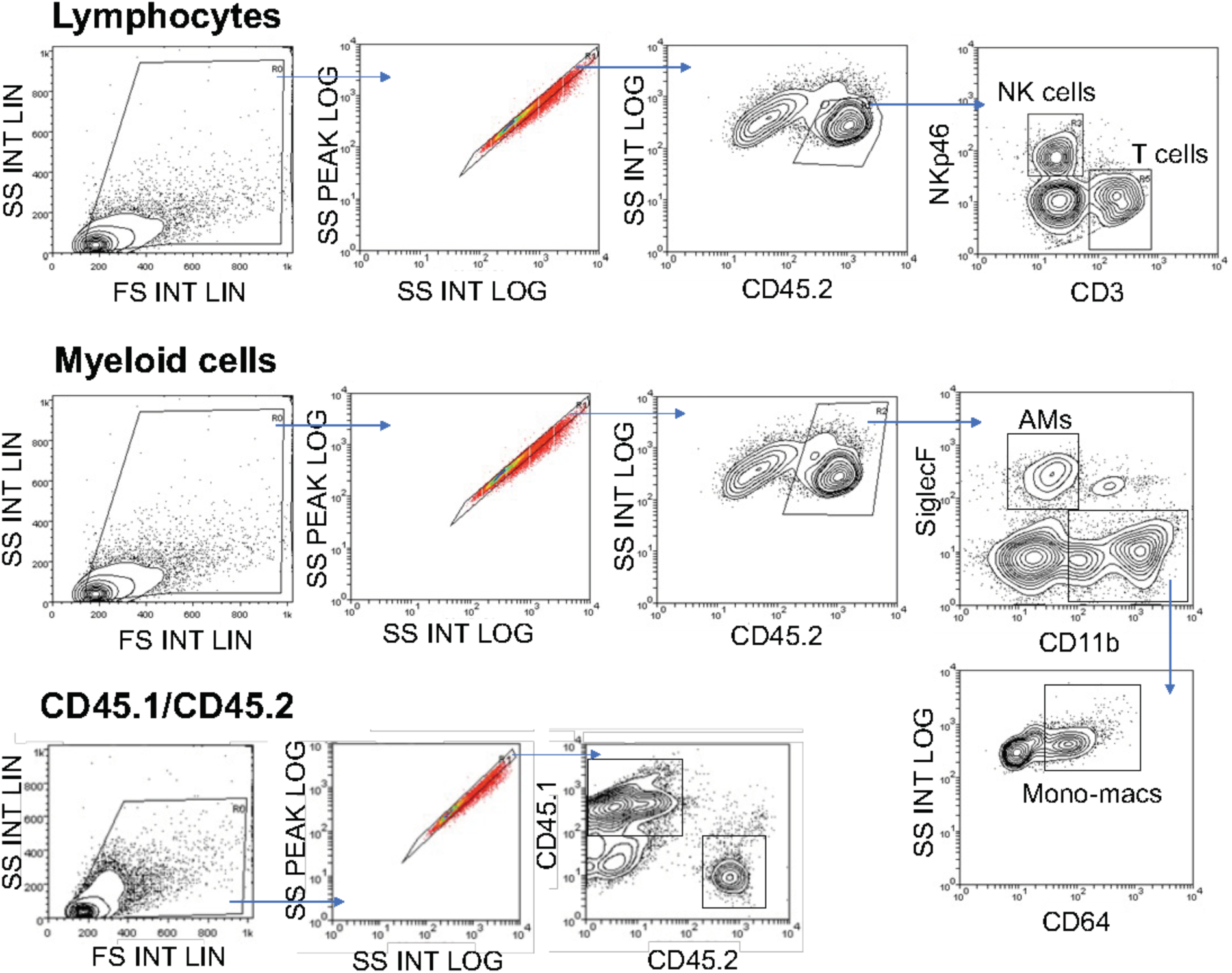
Gating strategies for immune cell subset identification by conventional flow cytometry.

**Supplementary Figure 9.**
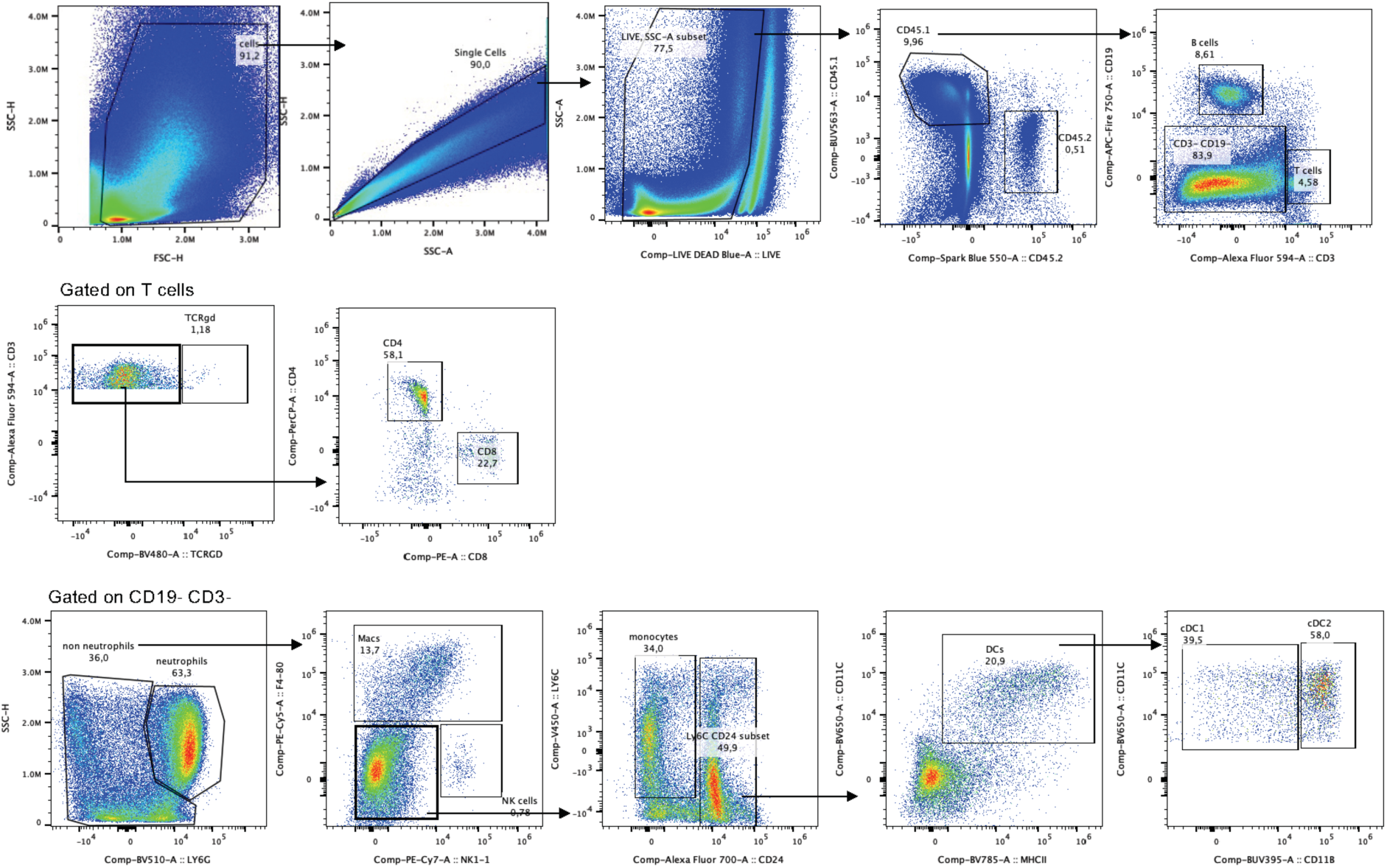
Gating strategies for immune cell subset identification by spectral flow cytometry (relevant to Fig. 6).

